# Genetically encoded live cell sensor for tyrosinated microtubules

**DOI:** 10.1101/2020.03.29.013250

**Authors:** Shubham Kesarwani, Prakash Lama, Anchal Chandra, P. Purushotam Reddy, AS Jijumon, Satish Bodakuntla, Balaji M Rao, Carsten Janke, Ranabir Das, Minhajuddin Sirajuddin

## Abstract

Microtubule cytoskeleton exists in various biochemical forms in different cells due to tubulin post-translational modification (PTMs). These PTMs are known to affect microtubule stability, dynamics and interaction with MAPs and motors in a specific manner, widely known as tubulin code hypothesis. At present there exist no tool that can specifically mark tubulin PTMs in live cells, thus severely limiting our understanding of tubulin PTMs. Using yeast display library, we identified a binder against terminal tyrosine of alpha tubulin, a unique PTM site. Extensive characterization validates the robustness and non-perturbing nature of our binder as tyrosination sensor, a live cell tubulin nanobody specific towards tyrosinated or unmodified microtubules. Using which, in real time we followed nocodazole, colchicine and vincristine induced depolymerization events of unmodified microtubules, and found each distinctly perturb microtubule polymer. Together, our work describes the tyrosination sensor and potential applications to study microtubule and PTM processes in living cells.

## Introduction

The cytoskeleton tubular polymer, microtubules performs diverse cellular functions, including but not limited to intracellular cargo transport, chromosome segregation and motility. These cellular processes involving microtubules are mediated by interactions with a cohort of molecular motors and microtubule associated proteins (MAPs). A key regulatory process that governs microtubule interaction with its cognate proteins is the diversity of tubulin genes and post-translational modifications (PTMs) (Janke, 2014). Most of the PTMs are reversible and defects in these PTM enzymes leads to abnormal levels of microtubule modifications, manifested in different disease pathologies (Magiera et al., 2018a) causing neurodegeneration (Magiera et al., 2018b) and cardiomyopathies (Chen et al., 2018; Robison et al., 2016).

Among the PTMs, tyrosination and detyrosination cycle of alpha tubulin carboxy-terminal site was the first PTM reported from rat brain extracts (Barra et al., 1973) and later in metazoans, ciliates and flagellates (Janke, 2014). The genetically encoded terminal tyrosine residue can be enzymatically removed by Vasohibin-SVBP complex, a detyrosinase identified recently (Aillaud et al., 2017; Nieuwenhuis et al., 2017). The tubulin tyrosine ligase (TTL), was the first tubulin PTM enzyme discovered, which reverses the detyrosination modification by adding tyrosine back to the terminal site of alpha tubulin (Barra et al., 1973). Over the years several additional tubulin modifications and respective enzymes have been identified across species; acetylation (L’Hernault and Rosenbaum, 1985), glutamylation (Eddé et al., 1990) and glycylation (Redeker et al., 1994). These tubulin modifications, except acetylation, occur at the carboxy-termini tails (CTT) of either alpha and/or beta tubulin gene products. The PTMs can also be combinatorial, overlapping with the diverse tubulin gene products creating diverse biochemical forms of microtubules across cell types (Janke, 2014), which makes tubulin PTM studies a challenging prospect to probe. Recent advances in protein engineering and expression have allowed creating homogenous microtubules with a particular PTM (Sirajuddin et al., 2014; Ti et al., 2018; Valenstein and Roll-Mecak, 2016; Vemu et al., 2014). These studies have highlighted that each PTM uniquely modulates different molecular motors (Barisic et al., 2015; McKenney et al., 2016; Sirajuddin et al., 2014), MAPs (Bonnet et al., 2001) and severing enzymes (Valenstein and Roll-Mecak, 2016), providing first insights into the regulatory roles of tubulin diversity.

A shortcoming of *in vitro* reconstitution experiments using homogenous modified microtubules is that it may not reflect the *in vivo* scenario, since microtubules inside cells can possess multiple PTMs at the same time. For example, the stable microtubules have been frequently associated with detyrosinated and acetylated microtubules (Bulinski et al., 1988). Similarly, the glutamylation and glycylation can occur at multiple sites of same tubulin CTTs and have been reported to co-exist in axonemal microtubules (Wloga et al., 2017). A typical cellular or *in vivo* study involving tubulin PTM involves genetic (Barisic et al., 2015; Magiera et al., 2018b), ectopic expression (van Dijk et al., 2007; Souphron et al., 2019) and chemical perturbations (Aillaud et al., 2017; Nieuwenhuis et al., 2017) of PTM enzymes, followed by antibody staining that are specific towards the respective PTM epitope (van Dijk et al., 2007; Gadadhar et al., 2017; Janke, 2014). Although the antibodies have illuminated the tubulin modifications, it severely limits our understanding of the spatial-temporal component of tubulin PTMs. Therefore, a cellular sensor which can detect and track tubulin modifications in real time will aid studying tubulin PTM dynamics and their function *in vivo*.

In general, the most common methods to label microtubules in living cells either involve fluorescent tagged alpha tubulin (Gierke et al., 2010; Kamath et al., 2010; Rusan et al., 2001), MAPs (Bulinski et al., 2001) or SiR-tubulin, a taxol derivative (Lukinavičius et al., 2014). The fluorescent protein tagged alpha tubulin can impede incorporation into microtubule polymer, this could be due to improper folding or incompatible tubulin isotypes. The fluorescent markers using MAPs and SiR-tubulin are known to affect microtubule assembly and/or dynamics. Nanobody and single chain antibody approaches have been successfully employed to study actin cytoskeleton and other PTMs (Helma et al., 2015), but without much success against microtubules (Traenkle and Rothbauer, 2017). So far two studies have attempted in this direction; nanobodies against microtubules, which was used to reconstruct super-resolution structures of microtubules (Mikhaylova et al., 2015). Another study has reported the identification of single chain antibody (anti-tubulin scFv) against tyrosinated form of microtubules (Cassimeris et al., 2013). However, the nanobody could not be employed in live cells (Mikhaylova et al., 2015) and no further study of anti-tubulin scFv application has been reported till date. Altogether there is a severe dearth of tools that can mark generic microtubules and/or tubulin PTMs in live cells.

To overcome this, we screened yeast display library (Gera et al., 2012) against CTT of alpha tubulin and identified a binder. Our study with the binder shows specificity against the tyrosinated state of microtubules, which does not interfere with the cellular or microtubule-based functions. The tyrosination sensor reported here, therefore becomes the first thoroughly characterized tubulin nanobody, that can be employed to follow unmodified (tyrosinated) microtubules in living cells.

## Results

### Strategy for screening binders against tyrosinated microtubule

Several studies have successfully employed carboxy-termini peptides of tubulins as epitopes to identify antibodies specific for a particular tubulin PTM (Bré et al., 1996; Gadadhar et al., 2017; Gundersen et al., 1984). Keeping this in mind, we synthesized the carboxy-terminus of TUBA1A (amino acids 440-451) with a biotin at the amino-terminus (biotin-TUBA1A 440-451). To obtain a binder protein specific for biotin-TUBA1A 440-451 (termed as *Hs*_TUBA1A), we employed a combinatorial yeast display library of SSO7d mutants screen as described previously (Gera et al., 2012). In order to select binders that are specific for the tyrosinated form, we also performed a negative selection of SSO7D library against biotin-TUBA1A 440-450 (detyrosinated, termed as *Hs*_TUBA1A-ΔY) and biotin-TUBA1A 440-451-[E]445 (mono-glutamylated, termed as *Hs*_TUBA1A-mG) peptides (Methods) (Figure 1A).

**Figure 1:**
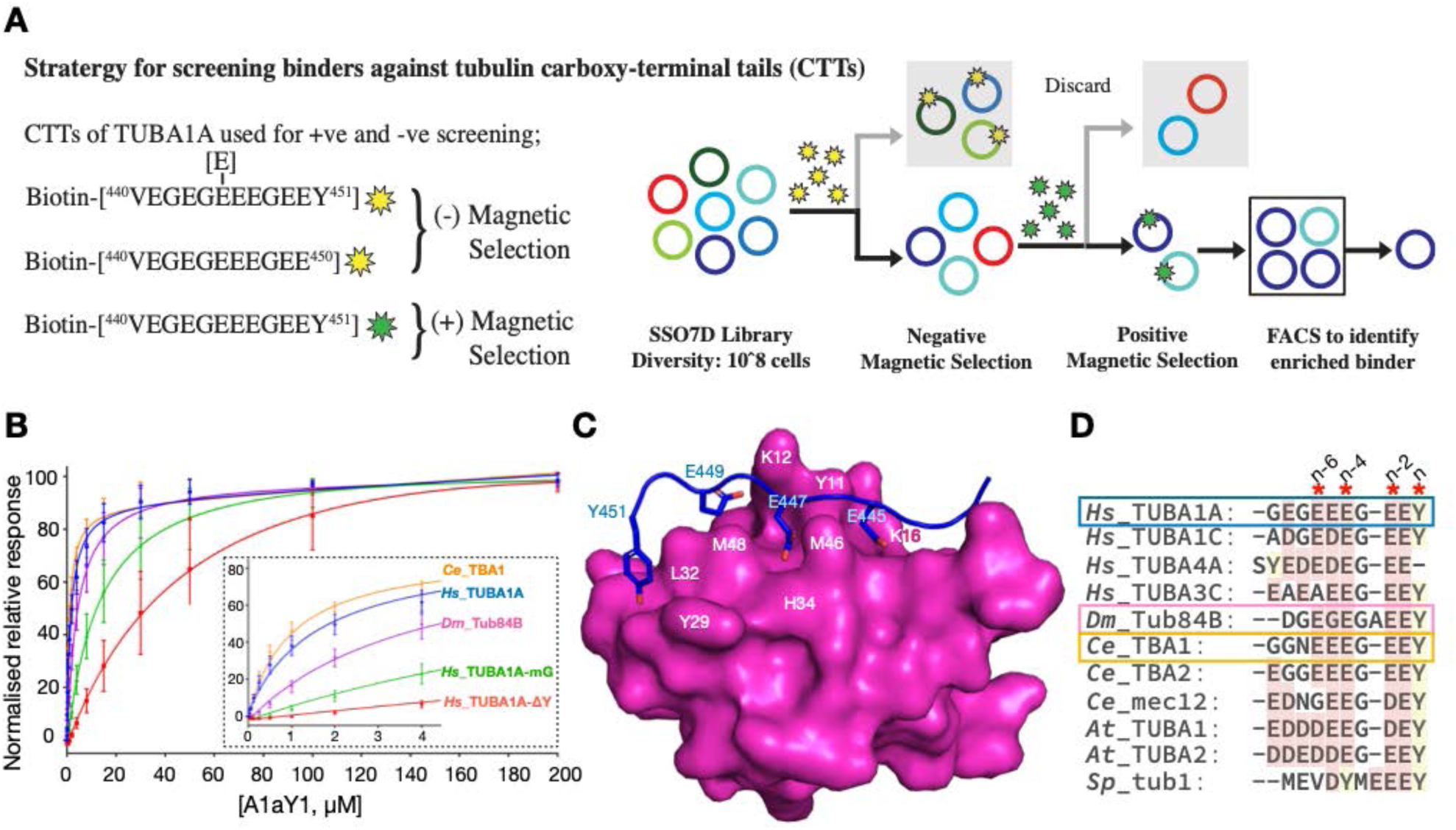
Identification, biochemical and structural characterization of A1aY1 binder. **A.** Schematic overview of the strategy employed to identify binder from SSO7D yeast display library. The biotinylated TUBA1A with Y (green star) and mono-Glu and ΔY (yellow star) used for positive and negative selection respectively. For detailed description see methods section. **B.** Dissociation constant (k_d_) of A1aY1 binder against biotinylated alpha tubulin CTT peptides of Human (*Hs*_TUBA1A; 1.6 ± 0.3μM, *Hs*_TUBA1A-ΔY; >60μM*, *Hs*_TUBA1A-mG; 13.6 ± 4.5μM), *Drosophila melanogaster* (*Dm*_Tub84B; 4.1 ± 0.9μM) and *C. elegans* (*Ce*_TBA1; 1.0 ± 0.2μM) measured using surface plasmon resonance state binding assay. Experiments were performed in triplicates with at least two different batches of the protein A1aY1 (* represents that the binding affinities can’t be uniquely determined with the current fit). The inset shows titration response up to 4μM A1aY1 binder concentration. **C.** The NMR structure of A1aY1 binder (magenta surface representation) bound to Hs_TUBA1A peptide (blue cartoon with key residues in stick representation). The key interacting residues from A1aY1 binder and alpha tubulin CTT are labeled in white and blue respectively. **D.** Sequence alignment of alpha tubulin CTTs from Human (*Hs*), *Drosophila melanogaster* (*Dm*), *Caenorhabditis elegans* (*Ce*), *Arabidopsis thaliana* (*At*), *Schizosaccharomyces pombe* (*Sp*) with gene names as indicated. The red asterisk indicates residues involved in A1aY1 binder interaction, the terminal tyrosine and alternating glutamate residues indicated as n series.

After four rounds of fluorescent activated cell sorting experiments (FACS) enrichment, 10 single yeast colonies were analyzed to identify the abundance of enriched clone(s) (Methods) (Supplementary Figure 1A). Among them two yeast clones A1aY1 and A1aY2 represented 30% and 20% enrichments respectively (Supplementary Figure 1B). Both these binders (A1aY1 and A1aY2) were purified as GFP tagged recombinant fusions and subjected to binding experiments with biotin-TUBA1A 440-451 peptide (Methods). The preliminary sensogram plots obtained from binding experiments strongly indicated that only A1aY1-GFP showed positive response towards biotin-TUBA1A 440-451 peptide (Methods) (Supplementary Figure 1C).

### Biochemical and structural characterization of the binder A1aY1

We further purified the A1aY-1 binder without any tags (Methods) and subjected to titration experiments with biotinylated TUBA1A 440-451, TUBA1A 440-450 and TUBA1A 440-451-[E]445 peptides, representing tyrosinated, detyrosinated and mono-glutamylated forms of TUBA1A carboxy-terminal tails (CTT) respectively (Methods). The binder A1aY1 binds with 1.6μM, >60μM and 13.6μM affinity with tyrosinated, detyrosinated and monoglutamylated TUBA1A CTT peptides respectively (Figure 1B). The biochemical data strongly indicates the importance of tyrosine at the TUBA1A CTT peptide for recognition by the A1aY1 binder. In addition, the titration data with monoglutamylated peptide suggests that in addition to terminal tyrosine, the A1aY1 binder might have interactions with the residues along the TUBA1A 440-451 CTT peptide.

To understand the interaction of TUBA1A 440-451 peptide (epitope) with A1aY1 binder better, we performed NMR experiments to gain three-dimensional structural information (Methods) (Table 1). The highest ranked ensemble structure of binder with epitope show the AlaY1 binding site overlap with the diversified regions of SSO7D protein (Figure 1C and Supplement Figure 2A, B, C, D & E). The key interacting residues from TUBA1A 440-451 peptide includes; Y451, E449, E447 and E445, the most common glutamylation site (Eddé et al., 1990) (Figure 1C and D). The terminal tyrosine (Y451, n^th^ residue) is latched with the aid of L32 and Y29 of A1aY1 binder, additionally the side chain hydroxy group of tyrosine makes a hydrogen bond with main chain amino group of G30 residue (Figure 1C and Supplement Figure 2). The remaining glutamic acid residues of TUBA1A 440-451 peptide alternatively make electrostatic contacts; E449 (n-2^nd^), E447 (n-4^th^) and E445 (n-6^th^) with K12, H34 and K16 of the A1aY1 binder respectively (Figure 1C).

**Table 1.**
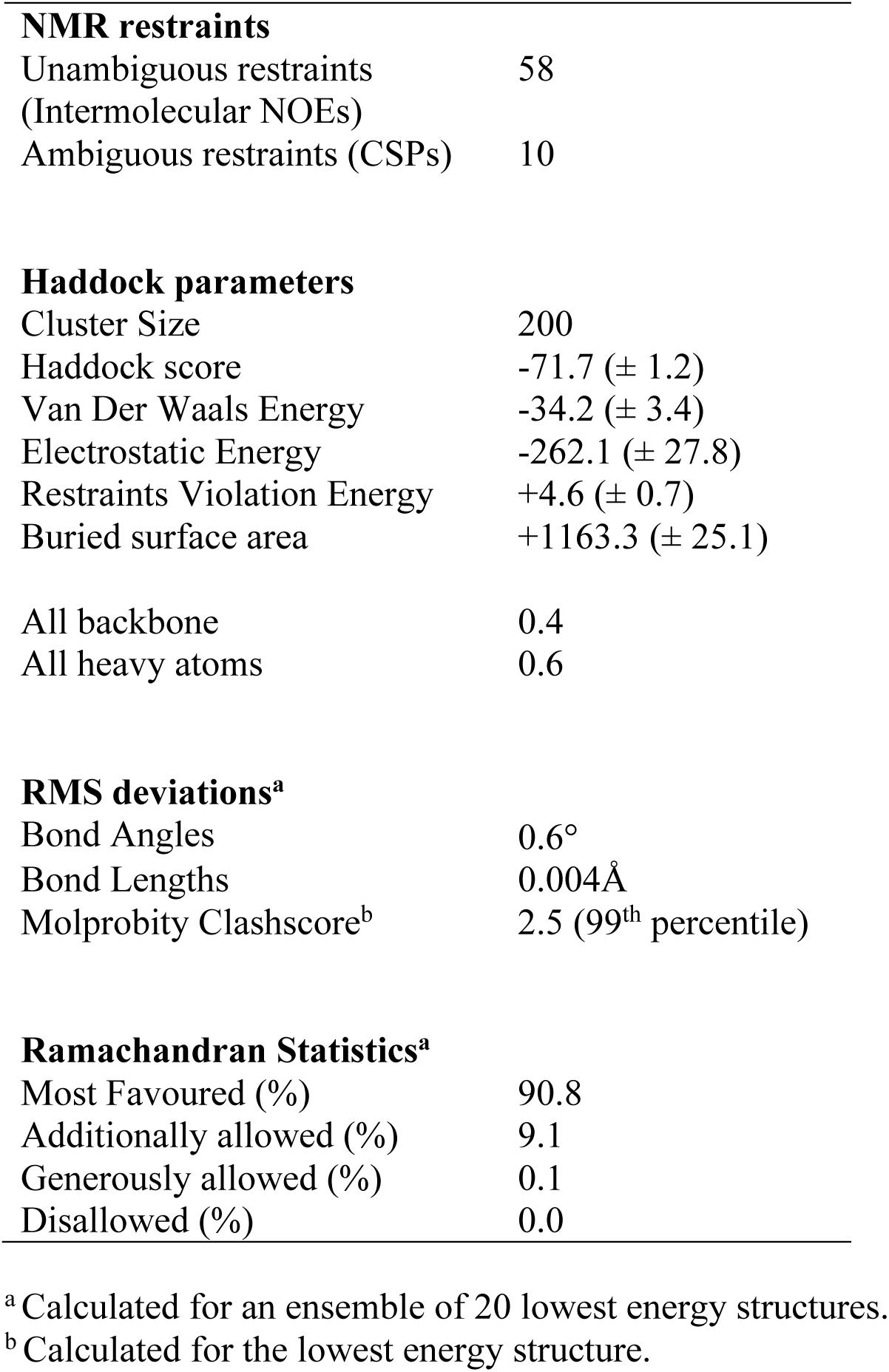
NMR and refinement statistics of the Binder/α-tubulin complex

Guided by the structure, we then compared ubiquitous sequences of alpha tubulin carboxy-termini from human, drosophila, worm, plant and fission yeast (Figure 1D). The terminal tyrosine (n^th^ residue) and the alternating glutamic acids (n-2^nd^, -4^th^, -6^th^ residues) are invariant across different species (Figure 1D). To further probe the CTTs interaction with binder, we tested *Dm*_Tub84B and *Ce*_TB1A of *D. melanogaster* and *C. elegans* alpha tubulin CTT peptides respectively, against our A1aY1 binder (Methods). Titration assays show that the *Dm*_Tub84B and *Ce*_TB1A CTTs interact with 1.0μM and 4.0μM affinities similar to the *Hs*_TUBA1A 440-451 CTT peptide (Figure 1B). In summary, the A1aY1 binder recognizes alpha tubulin CTTs from different organisms and the terminal tyrosine is an important element for A1aY1 binder interaction.

### A1aY1 binder labels specifically tyrosinated microtubules in cells

So far, our biochemical and structural experiments with A1aY1 binder have been limited to CTT peptide. In order to check whether A1aY1 binder can recognize microtubules in cells we generated a series of fluorescent protein fusion constructs and transiently expressed them in U2OS cells (Methods). Among them only TagBFP and TagRFP-T fused A1aY1 showed good colocalization with microtubules (Supplement Figure 3). Compared to the *Entacmaea quadricolor* fluorescent protein variants (TagBFP and TagRFP-T) the other fusion proteins showed poor signal to noise ratio and did not feature prominent microtubule signal (Supplement Figure 3). Both the TagBFP and TagRFP-T A1aY1 sensor illuminated similar microtubule features with amino- and carboxy-terminal fusions (Figure 2 and Supplement Figure 3). Therefore, we focused on TagBFP and TagRFP-T fused A1aY1 (hereafter, A1aY1 blue and red sensor respectively) and stably expressed them in U2OS cells (Figure 2) (Methods). In our stable cell lines, we also observed that upon high expression the microtubules begin to bundle in few cells (Supplement Figure 4). Thus, a medium to low-level expression of the blue and red A1aY1 sensor offers its application as a live cell biosensor for labelling microtubules (Figure 2 & Supplement Figure 4).

**Figure 2:**
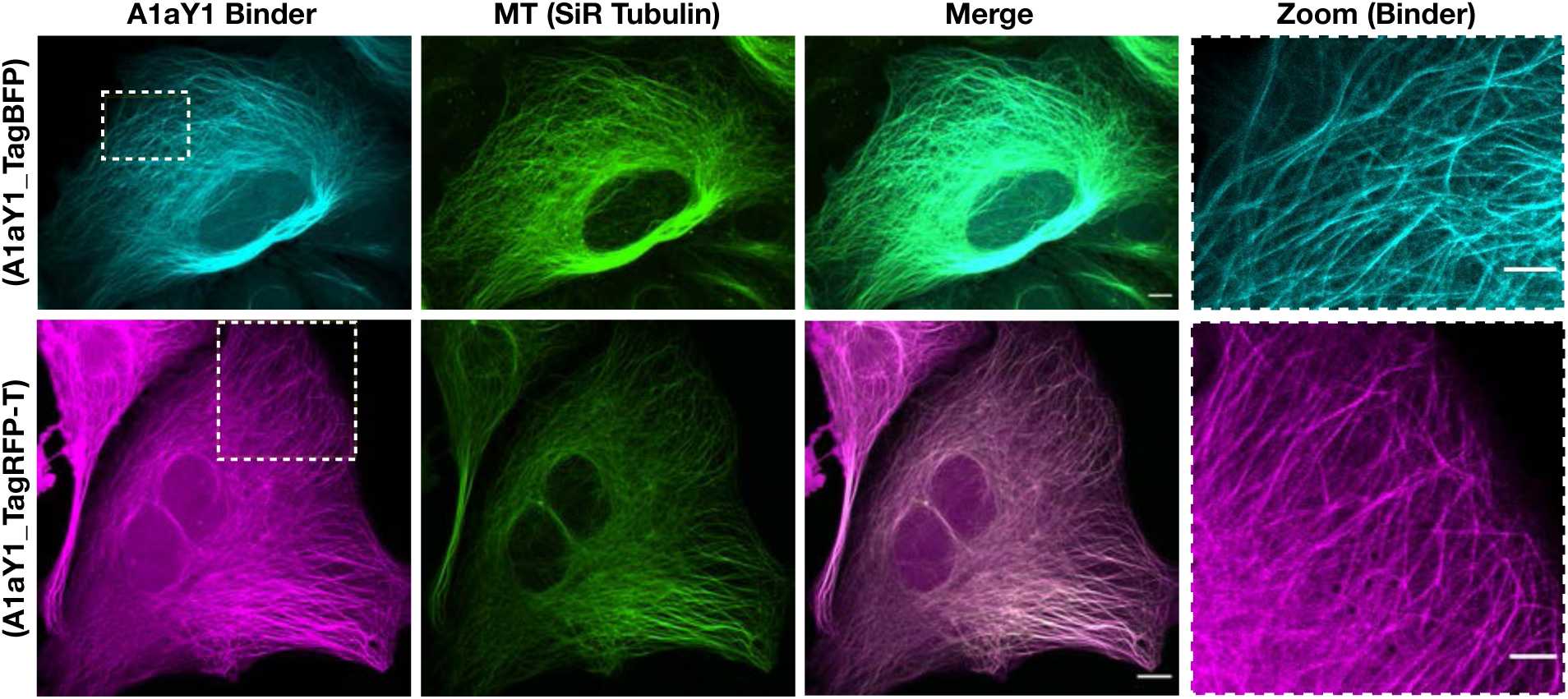
Fluorescent protein tagged A1aY1 binder labels cellular microtubules. Z-projection of confocal image stacks of U2OS cells stably expressing A1aY1 binder tagged with TagBFP or TagRFP-T at the amino terminus, in cyan and magenta respectively. The cells were additionally stained with SiR-tubulin (in green). The zoom panel shows closer view of microtubules labelled with A1aY1 TagBFP and TagRFP-T as indicated. Scale bar = 10μm and 5μm for the whole cell and zoomed panel respectively.

Tyrosinated microtubules are abundant at the interphase stage of fibroblasts and epithelial cells, therefore we inferred that our A1aY1 binder is marking bulk of the microtubules in our experiments. Our biochemical results show that A1aY1 binder is specific for tyrosinated peptide, we then checked the specificity of A1aY1 sensor towards detyrosinated and glutamylated microtubules. The U2OS stable cell lines of red A1aY1 sensor were co-transfected with detyrosinase (Aillaud et al., 2017; Nieuwenhuis et al., 2017) and polyglutamylation enzymes (Souphron et al., 2019) (Methods). For detyrosinase, we used VASH2_X1+SVBP enzyme complex, which shows elevated levels of detyrosinated microtubules upon expression in cells (Supplement Figure 5). TTLL5, its catalytically inactive version and TTLL7 enzymes were used for inducing polyglutamylation modifications in interphase cells (Souphron et al., 2019).

Co-transfection of detyrosinase in the stable lines with A1aY1 red sensor leads to complete loss of microtubule signal by A1aY1 sensor (Figure 3A & B). In the case of TTLL5 and its catalytically inactive version, we see a decrease and retention of microtubule signal by A1aY1 sensor respectively (Figure 3A & B). Conversely, TTLL7, an enzyme which is more specific towards beta tubulin CTT (van Dijk et al., 2007), shows no loss of colocalization by A1aY1 sensor towards microtubules (Figure 3A & B). This indicates that polyglutamylation modification at the alpha-tubulin CTT (E445 residue) will sterically interfere with A1aY1 sensor (Figure 3C, D & E). Thereby reducing the microtubule binding ability of A1aY1 sensor, which is similar to our observation with biochemistry using mono-glutamylated peptide (Figure 1B).

**Figure 3:**
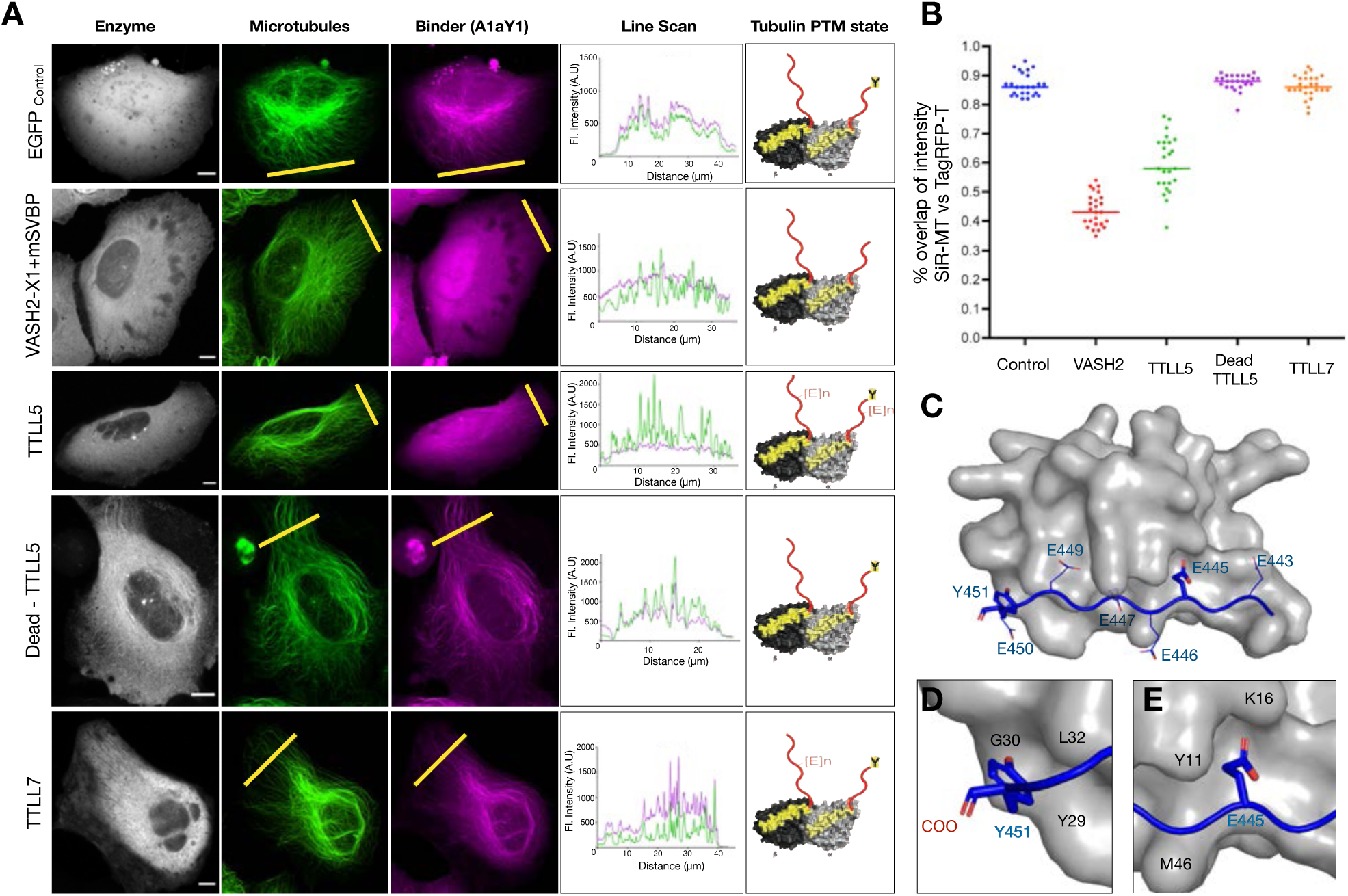
Specificity of A1aY1 binder towards tyrosinated microtubules. **A.** Confocal image stacks of U2OS cells stably expressing A1aY1 Tag-RFP-T, indicated as ‘Binder’ (magenta) along with GFP control, Vasohibin2-X1-GFP-2A-SVBP, TTLL5-eYFP, catalytically dead TTLL5-eYFP and TTLL7-eYFP, indicated as ‘Enzyme’ (grey) along with SiR-tubulin, indicated as ‘Microtubules’ (green). Line scans of SiR-tubulin (green) and A1aY1 Tag-RFP-T (magenta) fluorescence intensity signal for each panel as indicated by the yellow line. Cartoon representation of alpha/beta tubulin in right panel indicates the tubulin PTM state of microtubules for the respective PTM enzyme experiment. **B.** Quantification of Pearson’s coefficient (R-value) to calculate the correlation for the fraction of microtubules detected by A1aY1 Tag-RFP-T versus SiR-tubulin label, represented as ‘% overlap of intensity SiR-tubulin vs Tag-RFP-t’. A typical diffused cytoplasmic signal will have ∼ 50% or less overlap, whereas a complete co-localization will show near 100% overlap. n=25 cells, representing each dot for respective column. **C.** The NMR structure of A1aY1 (grey surface representation) and alpha tubulin CTT peptide (blue cartoon representation), the carboxy-terminal, tyrosine and glutamic acid residues are indicated. **D. and E.** Closer view of the terminal tyrosine (Y451) and polyglutamylation site glutamic acid (E445) with key interacting residues from A1aY1 binder as indicated.

In summary, our biochemistry and specificity experiments unequivocally suggest that A1aY1 sensor specifically recognizes tyrosinated microtubules. Therefore, hereafter we refer our A1aY1 binder as tyrosination sensor.

### Tyrosination sensor does not alter the cellular function of microtubules

Microtubules are majorly involved in mitotic spindle organization and chromosome segregation; in addition, a hallmark property of microtubules is their ability to undergo dynamic instability. Therefore, the next step in validation of tyrosination sensor as a live cell marker is to check whether they interfere with microtubule function. We first checked the viability and proliferative ability of stable U2OS cell lines expressing blue and red tyrosination sensor (Methods). Trypan blue assays, propidium iodide and DAPI based FACS for both blue and red sensor stable lines show that ∼95% of the cells are viable and in their proliferative state (Supplement Figure 6A & B). We also checked the ability of stable cell lines to undergo mitosis and chromosome segregation, here we could observe all the mitotic stages and image them using our tyrosination sensor (Supplement Figure 6A & B). This confirms that by stably integrating the sensor genes and constitutive expression of either blue or red tyrosine sensors does not interfere with cell viability and division.

Next, we measured microtubule dynamics in cells expressing tyrosination sensor using fluorescence from the TagRFP-T fused to A1aY1 binder (Methods). Time lapse images show that the microtubule polymerization events can be followed by TagRFP-T-A1aY1 fluorescence signal. Microtubules labelled with TagRFP-T-A1aY1 undergo typical dynamic instability states of growth, catastrophe and rescue (Figure 4A and Supplement Movie 1), suggesting that tyrosination sensor does not interfere with microtubule dynamics. In order to quantify the growth rates, we then transfected EB3-GFP and imaged microtubules along with EB3 comets (Methods) (Figure 4B, Supplement Figure 7A, B & C and Supplement Movie 2). We observed 0.35±0.09μm/s (n=450), 0.36±0.1μm/s (n=468) and 0.19±0.06μm/s (n=414) microtubule growth rates in the presence and absence of tyrosine sensor and with 0.5μM SiR-tubulin respectively (Figure 4C). Our live cell imaging with and without EB3-GFP comets also reveals that the tyrosination sensor binds to microtubules promptly during polymerization and disappears during catastrophe events (Figure 4A, B and Supplement Movie 1 & 2). Simultaneously, we measured EB3-GFP comets in the presence of SiR-tubulin, which shows the significant decrease in growth rates compared to the untreated and cells with tyrosination sensor (Supplement Figure 7). These live cell imaging experiments reveal that the tyrosination sensor can be used to follow microtubule polymerization and depolymerization events, without affecting their growth rates and dynamics (Supplement Movie 1 & 2).

**Figure 4:**
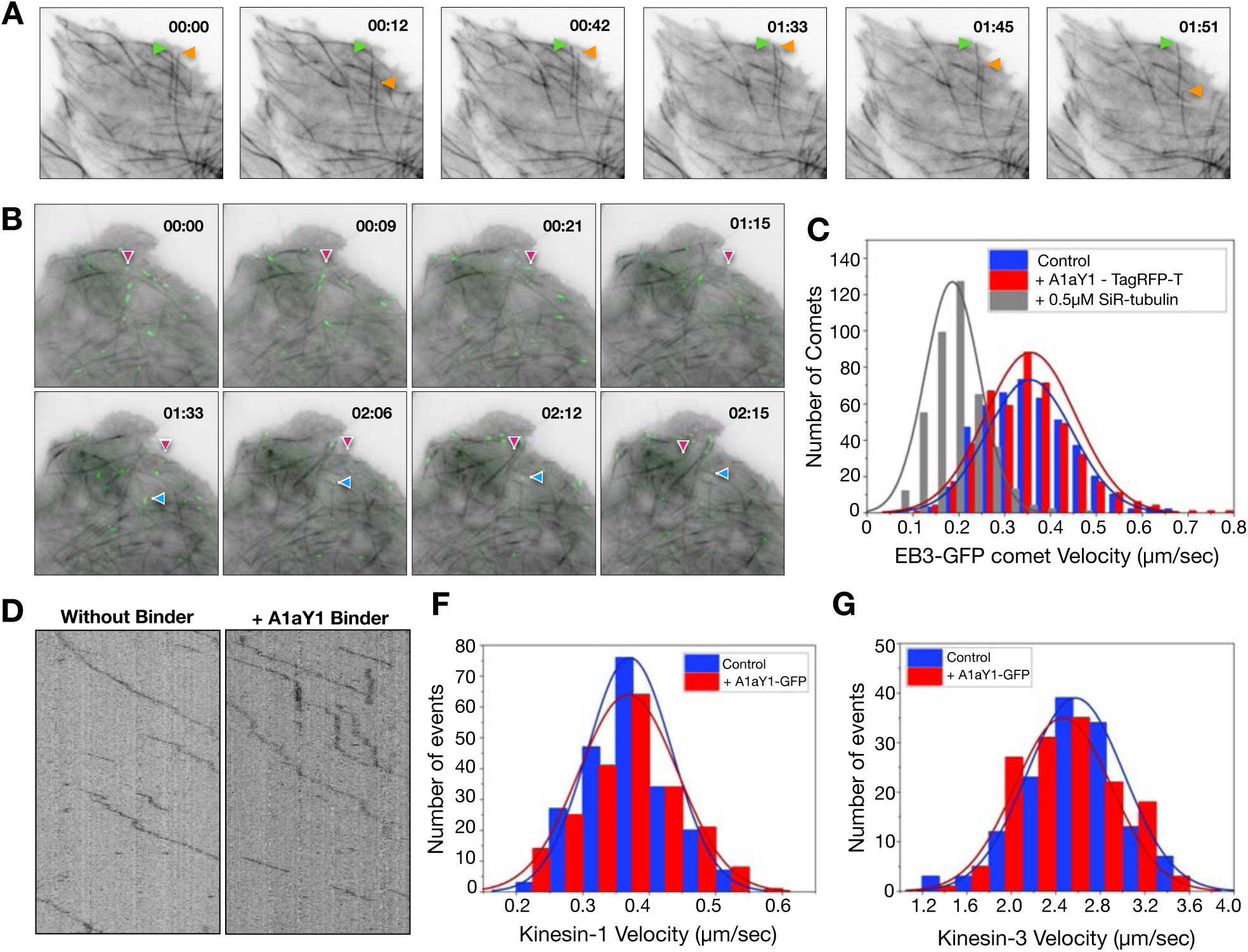
Effect of tyrosination sensor on microtubule function. **A.** Observation of microtubule dynamics using tyrosination sensor (A1aY1 Tag-RFP-T). The green arrow indicates microtubule growth pause and the orange arrow shows dynamic instability behaviour. Full movie of the frames can be found in Supplement movie 1. Scale bar = 5μm and time in minutes/seconds (mm:ss format) as indicated. **B.** Microtubule growth rate measurement using EB3 comet assay. Snapshots movie frames of stable U2OS cells with the red tyrosination sensor (in grayscale), transiently expressing EB3-GFP (in green). The magenta and blue arrows for reference of typical EB3 comets at microtubule plus-ends. Scale bar = 5μm and time format as indicated in A. **C.** Histogram of EB3 comet velocities, representing the microtubule plus-end growth rates (micron/sec). The average growth velocities for EB3 comets are 0.35±0.09μm/s (n=450), 0.36±0.1μm/s (n=468) and 0.19±0.06μm/s (n=414) for cells with tyrosination sensor (in red), without binder (in blue) and with 0.5μM SiR-tubulin (in grey), respectively. **D.** Kymograph of K560-SNAP motors moving on Hela microtubules with and without A1aY1 binder as indicated. Scale bar = 2μm and 25 seconds. **F. and G.** Histogram of K560-SNAP and KIF1A-1-393-LZ-SNAP motor velocities with (in red) and without (in blue) A1aY1 binder. The average velocity of K560-SNAP is 0.37± 0.06μm, n=214 (with A1aY1-GFP) and 0.37±0.07μm, n=208 (without A1aY1) and KIF1A-1-393-LZ-SNAP are 2.46±0.4μm, n=142 (with A1aY1-GFP) and 2.57±0.4μm, n=134 (without A1aY1). n represents number of motor particles analysed.

Another important microtubule function is their role as tracks for motor proteins, which facilitates intracellular cargo transport. In order to check if tyrosination sensor interferes with motor motility, we performed *in vitro* motility experiments using kinesin-1 (K560-SNAP) or kinesin-3 (KIF1A 1-393LZ-SNAP) and Hela microtubules (Souphron et al., 2019) marked with purified A1aY1-GFP as tyrosination sensor (Methods). Single molecule experiments show that both kinesin-1 and -3 does not show any major deviations from their normal motility behavior (Figure 4D, E & F and Supplement Figure 8).

Combinedly these experiments suggest that tyrosination sensor does not affect microtubule properties and their related cellular function, thus validating the suitability of our sensor for live cell experiments.

### Live cell imaging and mechanism of drugs that target microtubules

Currently live cell imaging of microtubules is carried out by fluorescent tagged alpha-tubulin, MAPs and SiR tubulin (fluorescent version of taxol) that stabilize microtubules (Supplement Figure 7). We envision that our tyrosination sensor, which does not affect dynamics (Figure 4) and specifically labels unmodified i.e., tyrosinated microtubules can be used to study microtubules in live cells. As a proof of concept, we employed drugs such as nocodazole, colchicine and vincristine that are known to target microtubules and are commonly used in cell biology studies to perturb microtubules. Although there are several reports about mode of drugs action, so far there are no reports of live cell imaging and description of depolymerization events in real time. To address this, we took advantage of our U20S cells stably expressing AlaY1-TagRFP-T, and were individually treated with either 10µM nocodazole, 500µM colchicine or 1µM vincristine (Methods). Upon nocodazole addition, we observed that the microtubules begin to shrink from the ends (Figure 5A, B & E, Supplement Movie 4), which is similar to the MCAK or kinesin-13 mediated end-depolymerization.

**Figure 5:**
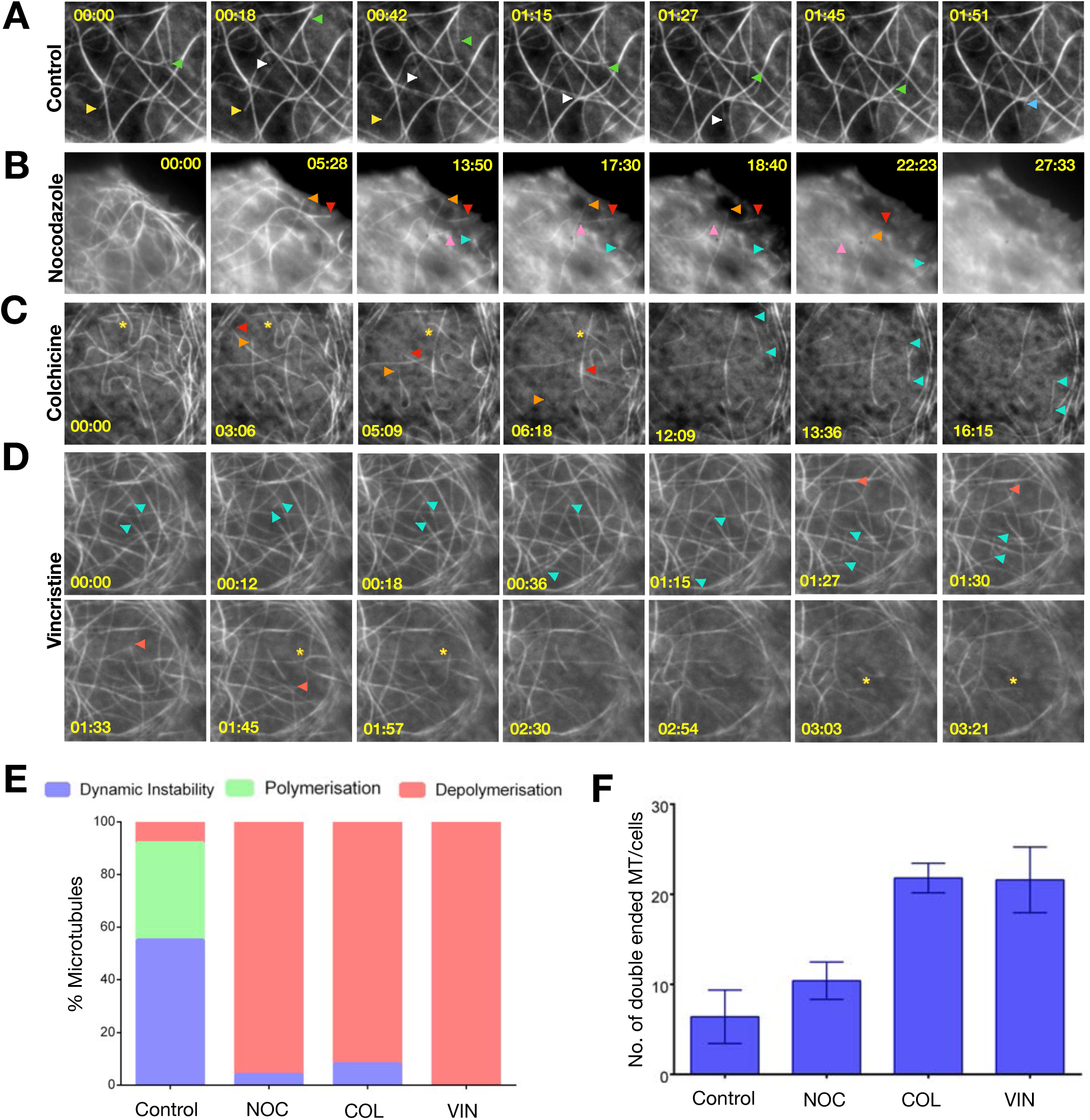
Drug induced microtubule depolymerization events in real time. **A.** Control (no drug) experiment of stable U2OS cells with red tyrosination sensor, microtubule dynamics are indicated with yellow, white, blue and green arrows. **B. C. and D.** Movie frames of stable U2OS cells with red tyrosination sensor treated with nocodazole, colchicine and vincristine (two rows). The depolymerization and severing events are indicated with coloured arrows and asterisks respectively for each panel. Full movie of the frames can be found in Supplement movie 3, 4, 5 and 6 for A, B, C, and D respectively. Scale bar = 5μm and time in minutes/seconds (mm:ss format) as indicated. **E.** Quantification of microtubule dynamics instability (in blue), where a catastrophe event is followed by growth, polymerization (in green) and depolymerization (in red) events respectively, from control and drug treated experiments. Data derived from 2-4 independent experiments n=100 events for each. **F.** The number of microtubules where both the ends are clearly visible, a proxy for microtubule severing events observed for control and drug treated experiments. Data derived from 2-4 independent batch of cells treated with drugs and averaged from 5 different cells; error bars represent SEM.

In the case of colchicine, we also observe that majority of the microtubules undergo end-on depolymerization events, with frequent severing like activity (Figure 5C, E & F, Supplement Movie 5). Similarly, we imaged vincristine mediated depolymerization events, vincristine when applied to cells the microtubules become brittle, similar to the filament severing activity (Figure 5D, E & F, Supplement Movie 6). Quantification of the depolymerization events by each drug shows that nocodazole, colchicine and vincristine have distinct mechanism of depolymerization as attributed by structural and biochemical studies (Gigant et al., 2005; Lee et al., 1980; Ravelli et al., 2004) (Figure 5E & F). While the mode of action for these drugs has been suggested earlier (Jordan and Kamath, 2007), here we are able to capture and follow the depolymerization events in real time. Thus, the A1aY1 sensor presents a great opportunity as a tool to study new microtubule targeting drugs and understand their mechanism in real time.

### Live cell super-resolution microscopy with tyrosination sensor

Cytoskeleton filaments and in general microtubules have been favorite test subjects for developing new methodology towards super-resolution imaging (Demmerle et al., 2015). Using the existing fluorescent tags, we performed 3color 3D-SIM on cells stably expressing tyrosination red sensor i.e., TagRFP-T-A1aY1 (Figure 6A) in interphase. Z-stacks were acquired for a total width of 2μm and all planes images were reconstructed, 3D volume rendered using alpha blending (Methods). Correlative analysis of SiR-tubulin versus TagRFP-T-A1aY1 signal in interphase cells shows a good agreement of colocalization (Pearson’s coefficient - 0.75; Spearman’s rank correlation – 0.88; Mander’s coefficient - 0.93) (Figure 6B), indicating the abundance of tyrosinated microtubules in during interphase cycle. Line scan comparison between 3D-SIM versus widefield filament width shows a gain of ∼300 nm in resolution (Figure 6C). Quantification of microtubules width for ∼50 filaments showed a mean FWHM value of 107±0.35 nm, in the region of half the diffraction limit as compared to the conventional resolution limit of ∼360 nm in a widefield setting (Figure 6D). We have also obtained 3D-SIM images for the mitotic stages in U2OS cells to confer the suitability of the probe for live cell super-resolution imaging over long periods of time (Supplement Movie 7).

**Figure 6:**
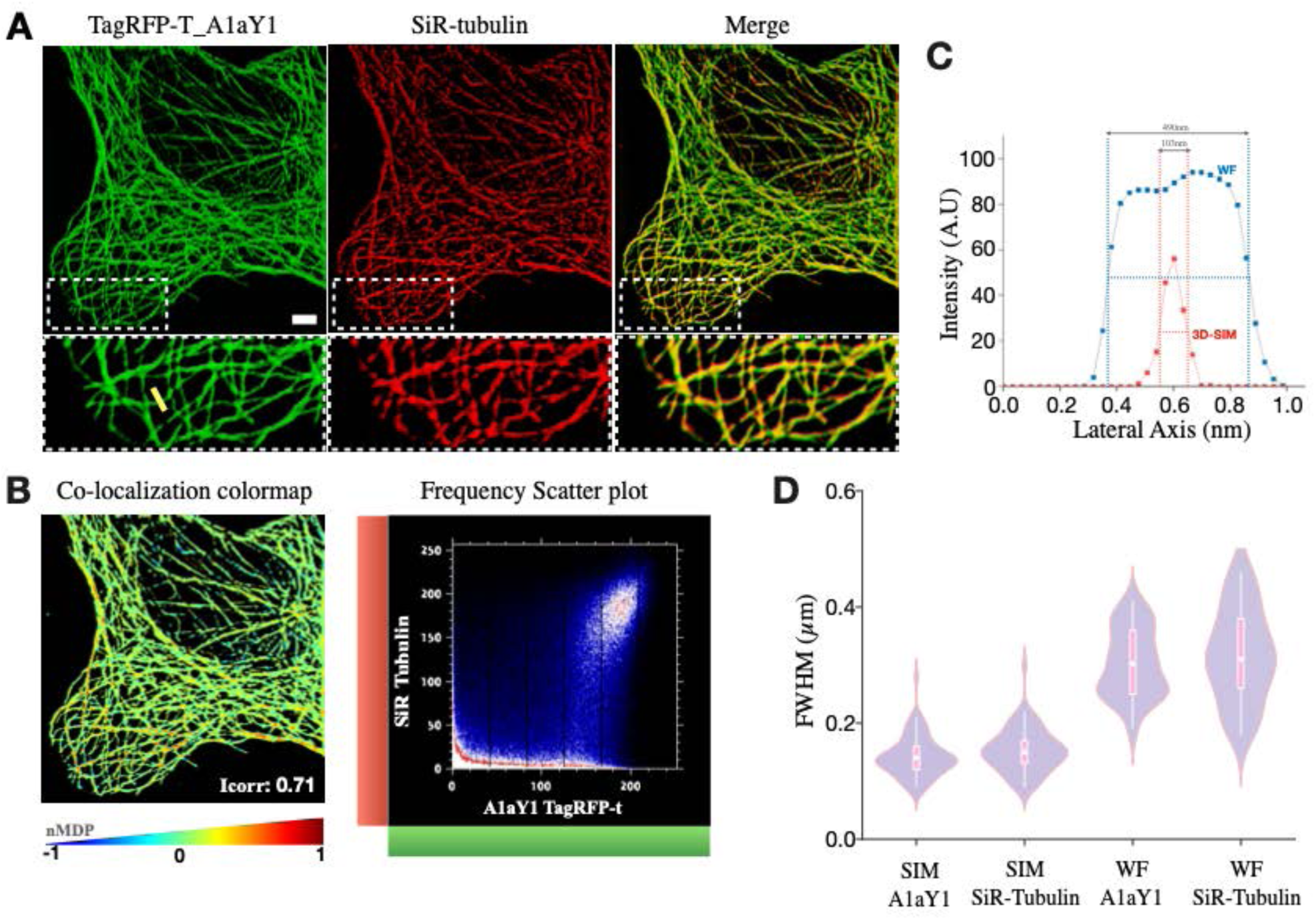
3D SIM imaging of microtubules using tyrosination sensor. **A.** Z-stack of 3D-SIM reconstructed images of amino-terminus TagRFP-T tagged to binder A1aY1 (red tyrosination sensor, shown in green here) stably expressed in interphase U2OS cells and labelled with SiR-Tubulin (red) (Scale bar = 2μm). The boxed areas are magnified below each panel. **B.** Colocalization colormap contains spatial distribution of calculated normalized mean deviation product (nMDP) values (ranging from -1 to 1). The distribution is based on a color scale in which negative nMDP values are represented by cold colors (segregation), values above 0 are represented by hot colors (colocalization). Representative frequency scatter plot for colocalization and pseudo-colormap of correlations between pairs of corresponding pixels in A1aY1 TagRFP-T and SiR-tubulin images, thereby offering quantitative visualization of colocalization. **C.** Line scans of representative single microtubules, from the position shown in the insets with yellow line, demonstrating the relative resolutions of 3D-SIM and widefield images. **D.** Quantification of FWHM measurements of n=50 microtubules for 3D-SIM and widefield images for both SiR-tubulin and red tyrosination sensor (TagRFP-T_A1aY1).

## Discussion

Fluorogenic nanobodies against intracellular structural components including actin cytoskeleton have been successfully employed in unraveling new biology. A notable exception has been the microtubule cytoskeleton (Traenkle and Rothbauer, 2017), in particular the tubulin PTMs where the epitopes are well-characterized and have yielded specific antibodies (Janke, 2014). In this study, we have employed alpha tubulin CTT peptide as epitope and discovered a binder from SSO7D library. The binder was then developed and validated as an intracellular nanobody against microtubules, specifically the tyrosinated form of alpha tubulin called tyrosination sensor. This sensor represents the first robust tubulin nanobody reported in the field, which does not affect microtubule and cellular functions. The imaging experiments with tyrosination sensor shows that single microtubule events can be followed in real time (Supplement Movie 1-6). Our EB3 comet assay also suggests that the tyrosination sensor does not interfere with binding of +TIP proteins, which contains CAP-Gly domains. We further extended the imaging capability of our tyrosination sensor towards super-resolution microscopy. Majority of the super-resolution studies with microtubules so far have used tubulin antibodies staining in fixed cells. Except in a few cases of live cell structural illumination microscopy (SIM), where the SiR-tubulin has been employed (Lukinavičius et al., 2014). Using SIM, here we demonstrate the application of tyrosination sensor in super-resolution microscopy, which is similar to the reported resolution of other SIM studies and comparable to the SiR-tubulin probe (Figure 6). However, the tyrosination sensor reported here does not affect microtubule dynamics compared to SiR-tubulin (Figure 4 C).

While the terminal tyrosine residue is an indispensable element in recognition by the sensor, our experiments indicates that the glutamic acid residues of alpha tubulin CTT also play important roles in tyrosination sensor binding. Tubulin CTTs contain a series of glutamic acid residues and provided that a terminal tyrosine is available, we predict that our tyrosination sensor will be able to bind to the tubulin/microtubule. This is strongly supported by our biochemical experiments with fly and worm alpha tubulin CTTs, which binds to the tyrosination sensor with similar affinities as that of human. A key element in this interaction is the third alternating glutamic acid residue (n-6^th^ residue) of alpha tubulin CTT, the most common site for glutamylation modification (Janke, 2014). Specificity experiments show that glutamylation modification at this site abolishes the interaction, suggesting that cross-linked glutamic acid residue will sterically hinder tyrosination sensor binding.

The alpha tubulin CTTs bearing terminal tyrosine are known to be recognized by CAP-Gly domains (Mishima et al., 2007), vasohibin or detyrosinase (Liao et al., 2019; Zhou et al., 2019), kinesin-13 and kinesin-2 motors (Sirajuddin et al., 2014). Additionally, the CAP-Gly domains bind to the carboxy-termini EEY motif of end-binding (EB) proteins, which is consensus to the alpha tubulin CTT terminus (Honnappa et al., 2006). Structural studies show that the sextette acidic motif (EEGEEY/F) of alpha tubulin CTT or the EEY motif of EBs are important for binding with CAP-Gly domain (Honnappa et al., 2006; Mishima et al., 2007). Vasohibin bound to alpha tubulin CTT complex structure also shows that the last five residues of alpha tubulin CTT (EGEEY) binds to the active site (Liao et al., 2019). In both cases the tyrosine recognition by CAP-Gly and VASH proteins reveals the importance of free mainchain carboxyl group of the terminal tyrosine residue. In contrast, our A1aY1:alpha tubulin CTT complex structure, reveals a unique mode of tyrosine sensing, which involves the interaction of phenyl ring and the hydroxyl group of tyrosine side chain (Figure 1C & 3D). Additionally, none of the naturally occurring proteins that bind to alpha tubulin CTT extend their recognition towards the glutamylation site (i.e., n-6^th^ residue). Therefore, we conclude that our tyrosination sensor senses the glutamylation state of alpha tubulin, in addition to the tyrosination state of microtubules.

Microtubules are well-known target for anti-cancer therapeutics, several reports describe drugs that stabilize and destabilize microtubules (Jordan and Kamath, 2007). Most of the drugs that are either used in therapeutics or for research purposes, have detailed account of their activity and binding site, from which the mechanism of action has been proposed. However, none of the work reported so far in this regard show drug induced microtubule depolymerization events. Here we applied our tyrosination sensor to image nocodazole, colchicine and vincristine induced microtubule depolymerization events in real time (Figure 5). Nocodazole is known to bind free tubulin dimers thus preventing their addition to microtubule polymer (Lee et al., 1980). Our results show that upon nocodazole treatment, the microtubules are in a constant state of catastrophe without any rescue events, in line with the proposed nocodazole mode of action. Colchicine was originally used to identify the tubulin (Borisy and Taylor, 1967), biochemical and structural investigations point that colchicine can bind to both soluble tubulin and as well as microtubule lattice (Jordan and Kamath, 2007; Ravelli et al., 2004). Cells when treated with colchicine show a combination of severed polymers and depolymerization events from both of the severed ends, underscoring its dual binding mode and twofold action. Vincristine is a potent anti-cancer drug, which binds only to microtubule polymer and destabilizes the lattice (Dhamodharan et al., 1995; Gigant et al., 2005; Jordan and Kamath, 2007; Jordan et al., 1992). In line with this structural finding, we observed more polymer severing and rapid depolymerization of double-ended microtubules.

The three drugs studied here show distinct modes of microtubule depolymerization, uniquely they also have commonalities amongst them. For example, nocodazole and colchicine when binds to the free tubulin traps them in curved state, which is incompatible for incorporation at the growing plus ends (Brouhard and Rice, 2014). Similarly, when colchicine or vincristine when binds to the microtubule, it induces lattice defects by kinking the longitudinal interaction between tubulins (Gigant et al., 2005; Ravelli et al., 2004). These lattice defects are then amplified leading to breaking of polymer, akin to severing like activity. In summary, the tyrosination sensor described here can be applied to study mechanisms pertaining to microtubule destabilizing drugs. In addition, since our tyrosination sensor marks unmodified microtubules without affecting dynamics, we predict that our sensor will be a valuable tool in screening new drugs that target microtubules. The specificity experiments described in Figure 3 outsets application towards studying PTM enzymes. The retention of sensor binding to microtubules upon the expression of catalytically dead TTLL5 suggests, that a similar approach will be advantageous in screening regulatory factors and drugs that affect PTM enzymes, such as vasohibin and TTLLs.

Together, the tyrosination sensor described here presents an opportunity to study microtubules and tubulin PTMs in live cells using fluorescence and super-resolution microscopy methods. Further providing a prospect to expand this methodology to generate sensors against other microtubule PTMs such as detyrosination, acetylation, glutamylation and glycylation.

## Materials and Methods

Detailed materials methods can be found in Supplement Information.

---

**Supplement Movie 1:** Visualization of cellular microtubules and dynamics using tyrosination sensor. Scale bar = 5μm and time in minutes/seconds (mm:ss format) as indicated.

**Supplement Movie 2:** Microtubule growth rate measurements using EB3-GFP (green) with tyrosination sensor (grey). Scale bar = 5μm and time in minutes/seconds (mm:ss format) as indicated.

**Supplement Movie 3:** Control (no drug) treated U2OS cells stably expressing red tyrosination sensor. The yellow box indicates the region represented in Figure 5A. Scale bar = 5μm and time in minutes/seconds (mm:ss format) as indicated.

**Supplement Movie 4:** 10μm Nocodazole treated U2OS cells stably expressing red tyrosination sensor. Scale bar = 5μm and time in minutes/seconds (mm:ss format) as indicated.

**Supplement Movie 5:** 0.5mM Colchicine treated U2OS cells stably expressing red tyrosination sensor. Scale bar = 5μm and time in minutes/seconds (mm:ss format) as indicated.

**Supplement Movie 6:** 1μm Vincristine treated U2OS cells stably expressing red tyrosination sensor. Scale bar = 5μm and time in minutes/seconds (mm:ss format) as indicated.

**Supplement Movie 7:** 3D-SIM reconstructions of interphase and mitotic phase of U2OS cells stably expressing red tyrosination sensor.

## Acknowledgements

The authors acknowledge the NMR facility of Biophysics Core and the Central Imaging and Flow Facility (CIFF) at NCBS/inStem/CCAMP campus. S.K and P.L supported by inStem Graduate Program. AC is a Wellcome-DBT ECF IA/E/15/1/502339. CJ is supported by the Institut Curie, the French National Research Agency (ANR) award ANR-17-CE13-0021, the Institut National du Cancer (INCA) grant 2014-PL BIO-11-ICR-1, and the Fondation pour la Recherche Medicale (FRM) grant DEQ20170336756. SB was supported by the FRM grant FDT201805005465. JAS was supported by the European Union’s Horizon 2020 research and innovation programme under the Marie Skłodowska-Curie grant agreement No 675737, and the FRM grant FDT201904008210. The NMR facility is supported by B-life grant from DBT(BT/PR5081/INF/156/2012). The project was partly supported by Tata Institute of Fundamental Research. R.D. acknowledges the DBT-Ramalingaswamy fellowship (BT/HRD/23/02/2006). inStem core grants from Department of Biotechnology, India, Wellcome Trust-DBT India Alliance Intermediate Fellow (IA/I/14/2/501533) and EMBO Young Investigator award to MS. C.J and M.S are jointly funded by CEFIPRA (5703-1).

## Author Contributions

SK and MS conceived the project. SK screened, validated and performed the microscopy experiments with binder. PL carried out the motor motility experiments. AC and SK performed 3D-SIM imaging. SB, JAS and CJ provided reagents for PTM specificity work and Hela tubulin. PPR and RD determined the NMR structure. BMR provided the yeast display library and valuable inputs in screening. MS supervised the project. SK and MS wrote the paper and all authors commented on the manuscript.

## Data Availability

NMR structure coordinates for A1aY1 binder: Alpha tubulin CTT complex are deposited in PDB under following code; PDB XXXX.

---

**Supplementary Figure 1:**
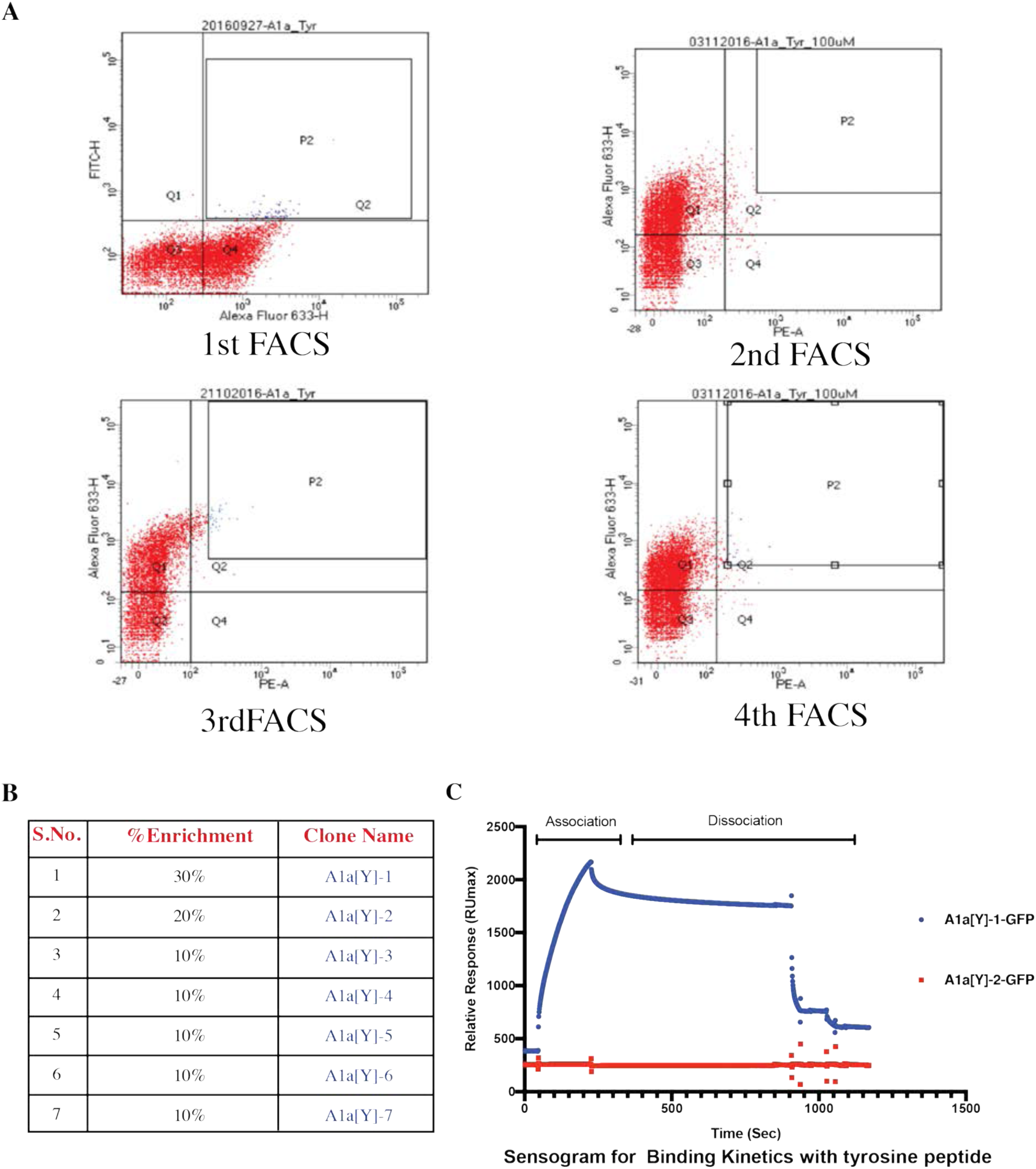
**A.** Representative images for the 10000 yeast cells sorted on BD FACS Aria fusion cell sorter. Total 4 rounds of sorting experiments were performed. The population P2 is marked in blue (Q2 quadrant) is collected (>4000 cells during each sorting). Q3 quadrant represents the unstained population. Expression of the binders in SSO7D library was marked by labeling the c-myc tag in Alexa fluor-633 channel (Q4 in 1st FACS and Q3 in 2nd, 3rd, 4th, FACS) and the binder bound peptide were labeled with FITC (1st FACS) or Phycoerythrin (PE, for 2nd, 3rd and 4th FACS) channel (see Methods). **B.** Table represents the abundance (% enrichment) of the clones obtained post 4th sorting after analyzing 10 single colonies. **C.** Sensogram obtained from the immobilized biotin-Hs_TUBA1A peptide on a streptavidin surface in the surface plasmon resonance experiment to determine the binding parameters of the two abundant clones (A1aY1 and A1aY2) for their binding parameters. ∼ 8μM binders (A1aY1 and A1aY2) were purified with GFP tag at carboxy-terminus.

**Supplementary Figure 2:**
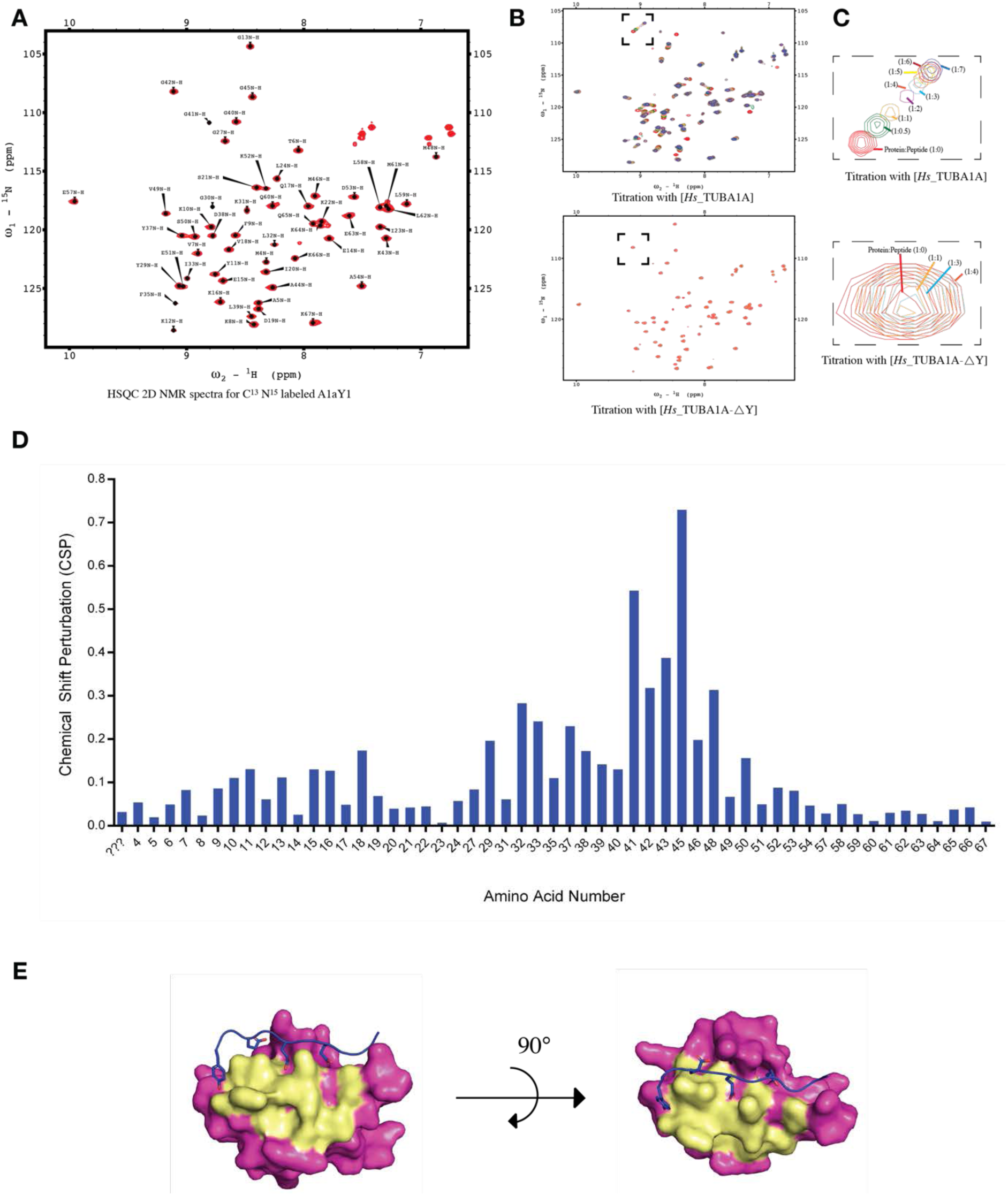
**NMR based structural determination and titrations with the alpha CTT peptides. A.** 2D NMR spectra (HSQC, Heteronuclear Single Quantum Coherence spectra) and assignment of the peaks for corresponding amino acids in A1aY1 protein labelled with ^15^N and ^13^C isotopes. The X-axis of the HSQC spectra represents the proton (^1^H) shift and Y-axis for ^15^N shift. **B.** HSQC spectra of the ^13^C ^15^N labelled A1aY1 protein in presence of alpha tubulin CTT peptides (*Hs*_TUBA1A and *Hs*_TUBA1A-ΔY). A titration of A1aY1 protein (200μM) with 1:0.5, 1:1, 1:2, 1:3, 1:4, 1:5, 1:6 and 1:7 excess of *Hs*_TUBA1A peptides showed positive chemical shift in the residues of A1aY1; while titration with *Hs*_TUBA1A-ΔY in 1:1, 1:2, 1:3 and 1:4 excess doesn’t show any significant change in the position of the residues. **C.** Zoomed in residue from the titration of *Hs*_TUBA1A and Hs_TUBA1A-ΔY (marked with black dashed box). **D.** Values of the chemical shift perturbations calculated from the ppm shift in the residues upon titration with *Hs*_TUBA1A. **E.** The NMR structure of A1aY1 binder (magenta) showing the residues randomized (yellow) in SSO7D structure lying in the binding pocket for *Hs*_TUBA1A peptide.

**Supplementary Figure 3:**
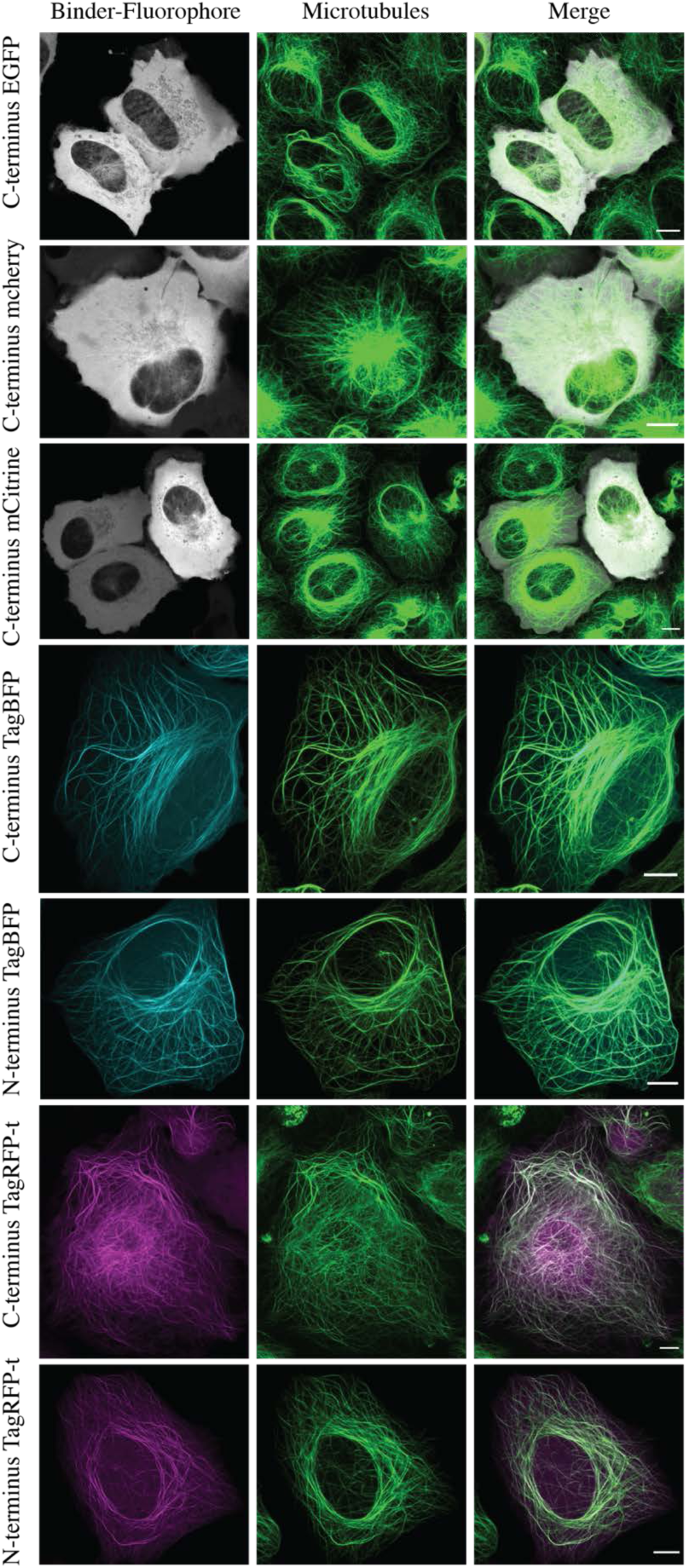
Transient transfection in wild type U2OS cells with carboxy-terminus eGFP (grey), mcherry (grey), mCitrine (grey), TagBFP (cyan) and TagRFP-t (magenta). Amino-terminus TagBFP (cyan) and TagRFP-T (magenta) fluorophore tagged A1aY1 binder. Total cellular microtubules were labeled with SiR-tubulin (shown in green). Scale bar = 10μm.

**Supplementary Figure 4:**
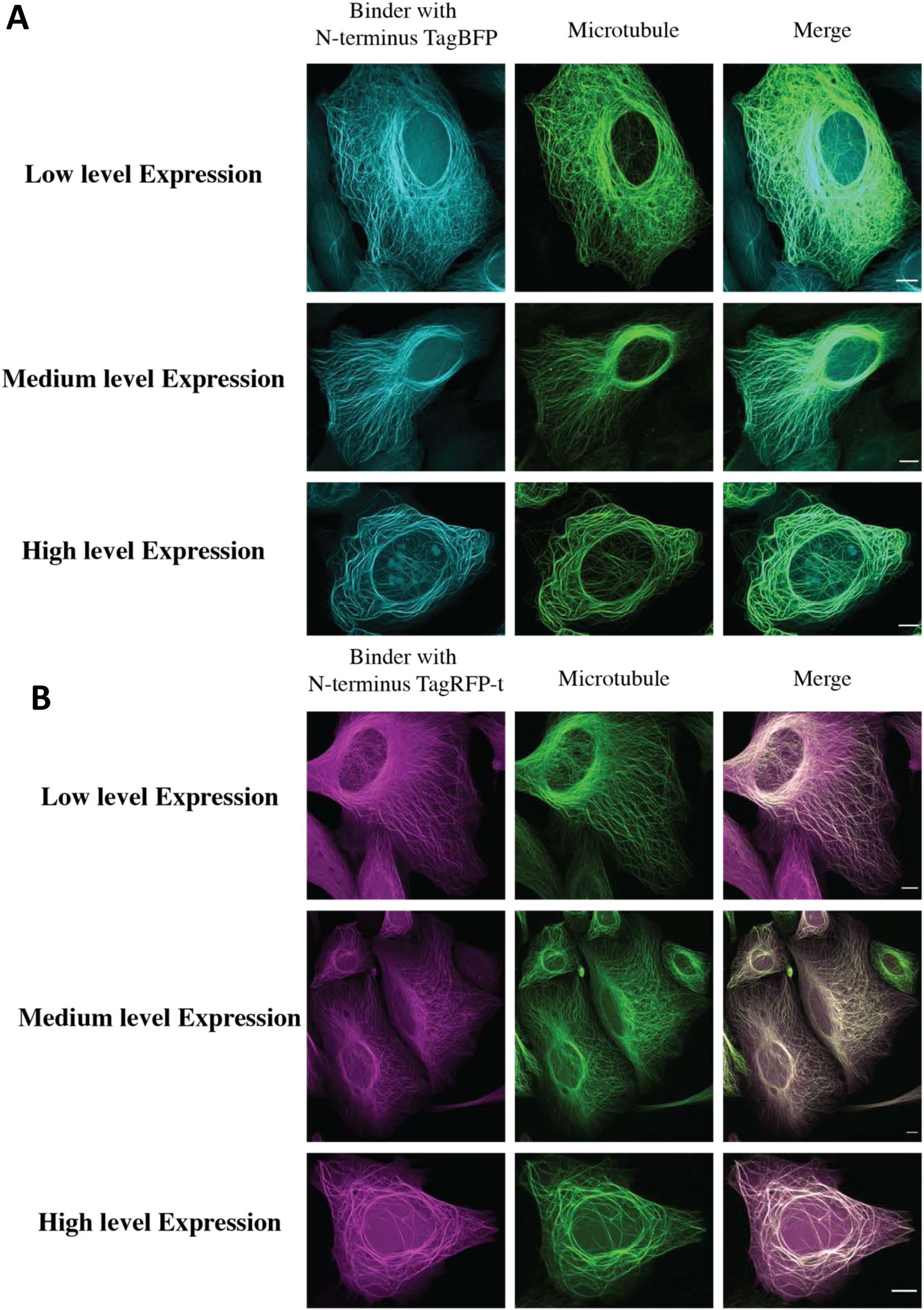
Differential expression of the A1aY1 binder stably incorporated in U2OS cells. **A.** Blue sensor (A1aY1-TagBFP) or **B.** Red sensor (A1aY1-TagRFP-T). High level expression of either blue or a red sensor leads to microtubule bundling. Low to medium level expression is optimal for A1aY1 application as a live sensor for tyrosinated microtubules. Scale bar = 10μm.

**Supplementary Figure 5:**
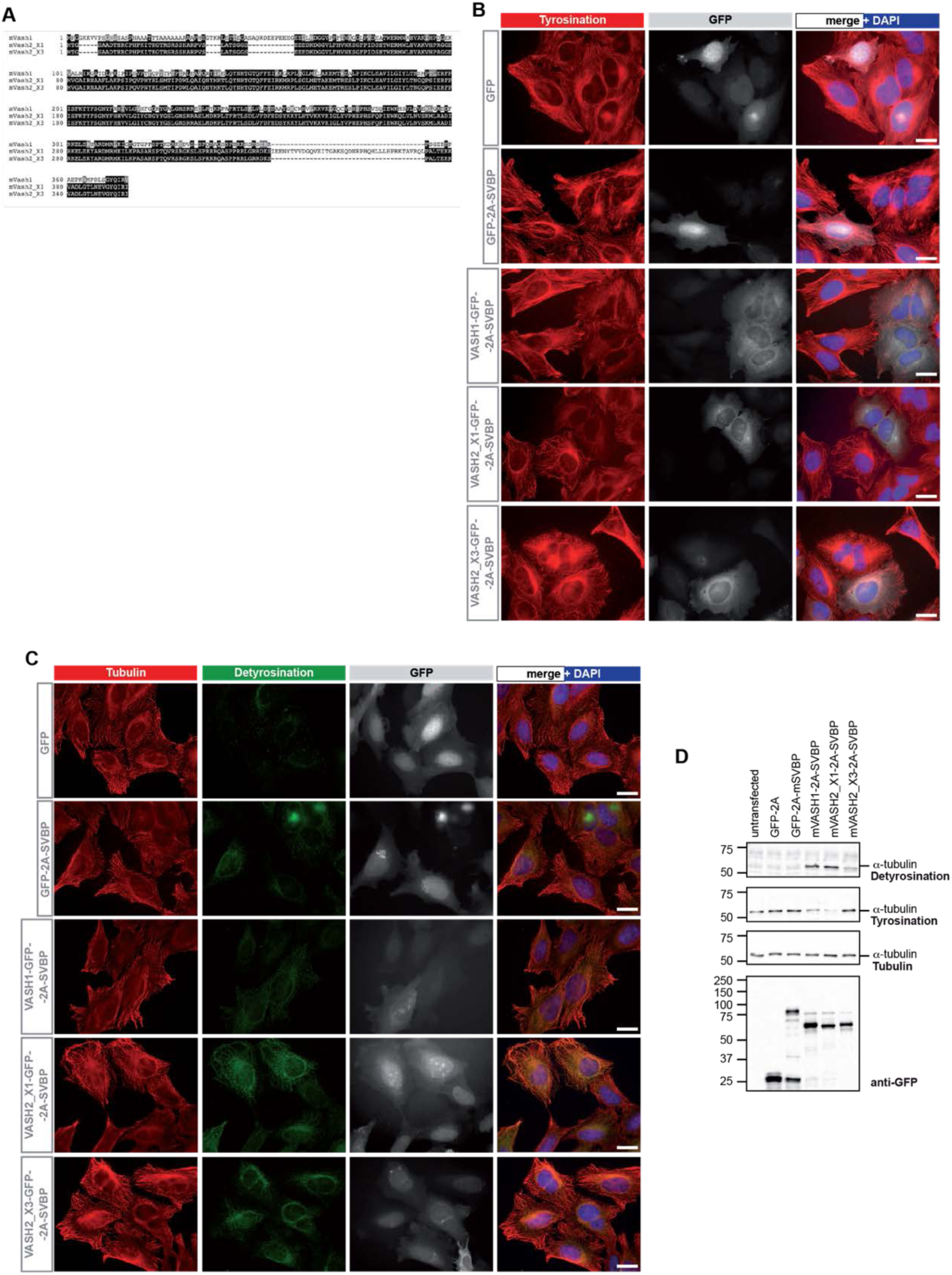
**A.** Sequence alignment of mVash1 and splice isoforms of mVash2 (mVash2_X1 and mVash2_X3) was shown. Identical amino acids across these proteins were highlighted in black and amino acids with similar charge were highlighted in grey color. mVash2_X1 contains an extra stretch of aminoacid sequence in the N-terminal region. **B.** HeLa cells transduced with lentivirus encoding different constructs of mVash-mSvbp, namely GFP-2A-mSVBP, mVash1-GFP-2A-mSVBP, mVash2_X1-GFP-2A-mSVBP, mVash2_X3-GFP-2A-mSVBP and GFP-alone were fixed and stained with tyrosination antibody (shown in red color). GFP expression and nuclear staining (DAPI) were shown in grey and blue colors, respectively. Of all the Vash-constructs, mVash1 and mVash2_X1 have shown a clear reduction in the microtubule tyrosination levels. Scale bar: 20 µm. **C.** HeLa cells transduced with lentivirus encoding different constructs of mVash-mSvbp, namely GFP-2A-mSVBP, mVash1-GFP-2A-mSVBP, mVash2_X1-GFP-2A-mSVBP, mVash2_X3-GFP-2A-mSVBP and GFP-alone were fixed, stained with antibodies against tubulin (shown in red color) and detyrosination (shown in green color). GFP expression and nuclear staining (DAPI) were shown in grey and blue colors, respectively. Note that in contrast to mVash1 and mVash2_X3, most of the microtubules in cells expressing mVash2_X1 are detyrosinated. Scale bar: 20 µm. **D.** Immuno blot analyses of HeLa cell lysates transduced with lentivirus encoding different constructs of mVash-mSVBP. Blots were probed with antibodies against detyrosination, tyrosination, α-tubulin and with anti-GFP for visualizing the expression of GFP-tagged mVash-proteins. Note the increase of detyrosination and decrease of tyrosination in the extracts expressing mVash1 and mVash2_X1.

**Supplementary Figure 6:**
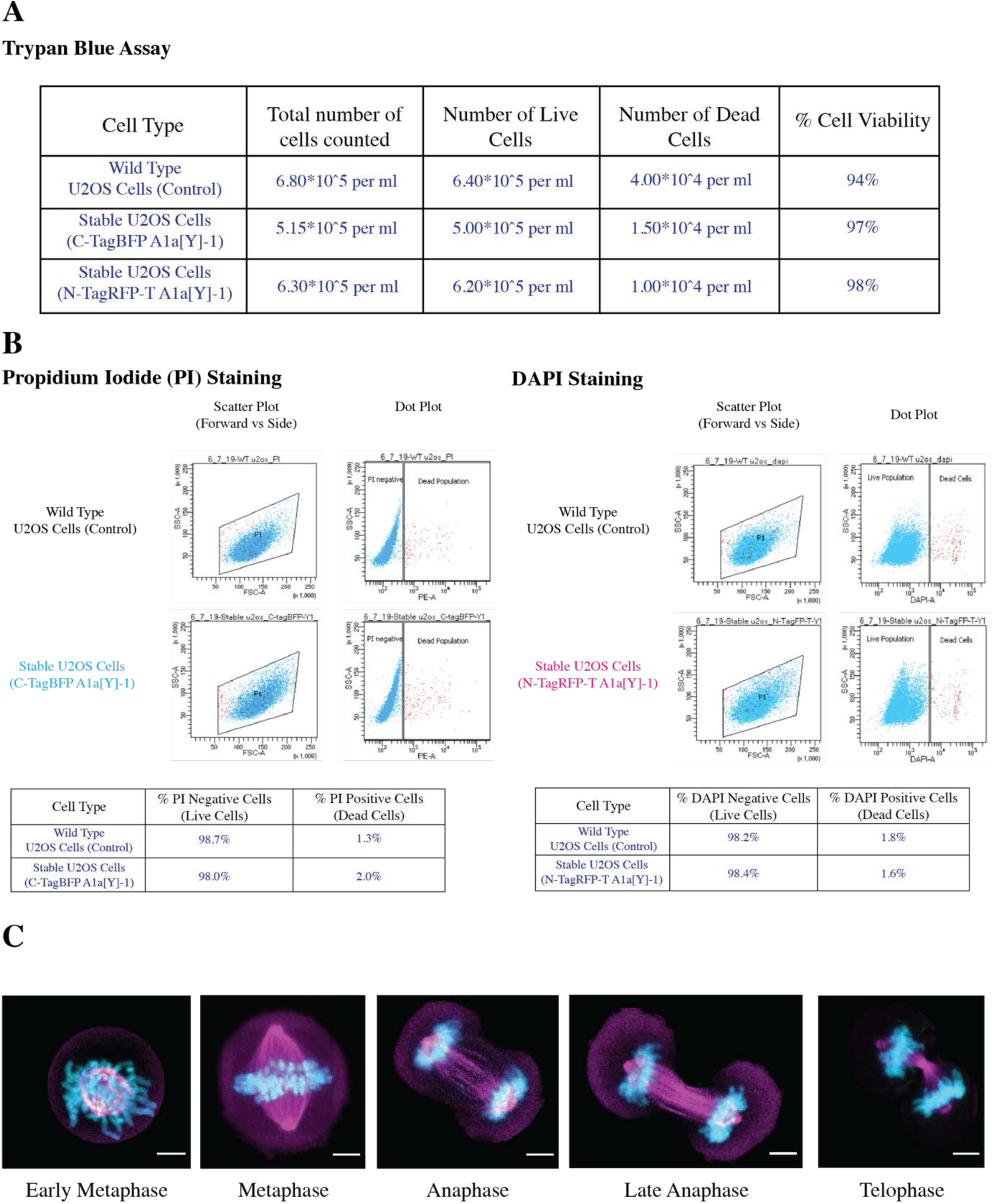
**Cell viability assays**. **A.** Trypan blue staining of the wild type U2OS cells and stable U2OS cells expressing blue sensor or the red sensor. In all the cases the cells were viable above 90%. **B.** Fluorescence cytometry-based viability test for red sensor expressing U2OS cells stained DAPI and blue sensor expressing U2OS cells stained with Propidium Iodide (PI). In both the cases the percentage of dead cells (DAPI positive or PI positive cells) doesn’t exceed more than 2% which is comparable to the wild type U2OS cells. **C.** U2OS cells stably expressing red sensor undergoes mitosis with cells imaged for different stages of cell cycle (metaphase, anaphase and telophase). Binder bound microtubules shown in magenta and the DNA marked with DAPI shown in cyan. Scale bar = 5μm.

**Supplementary Figure 7:**
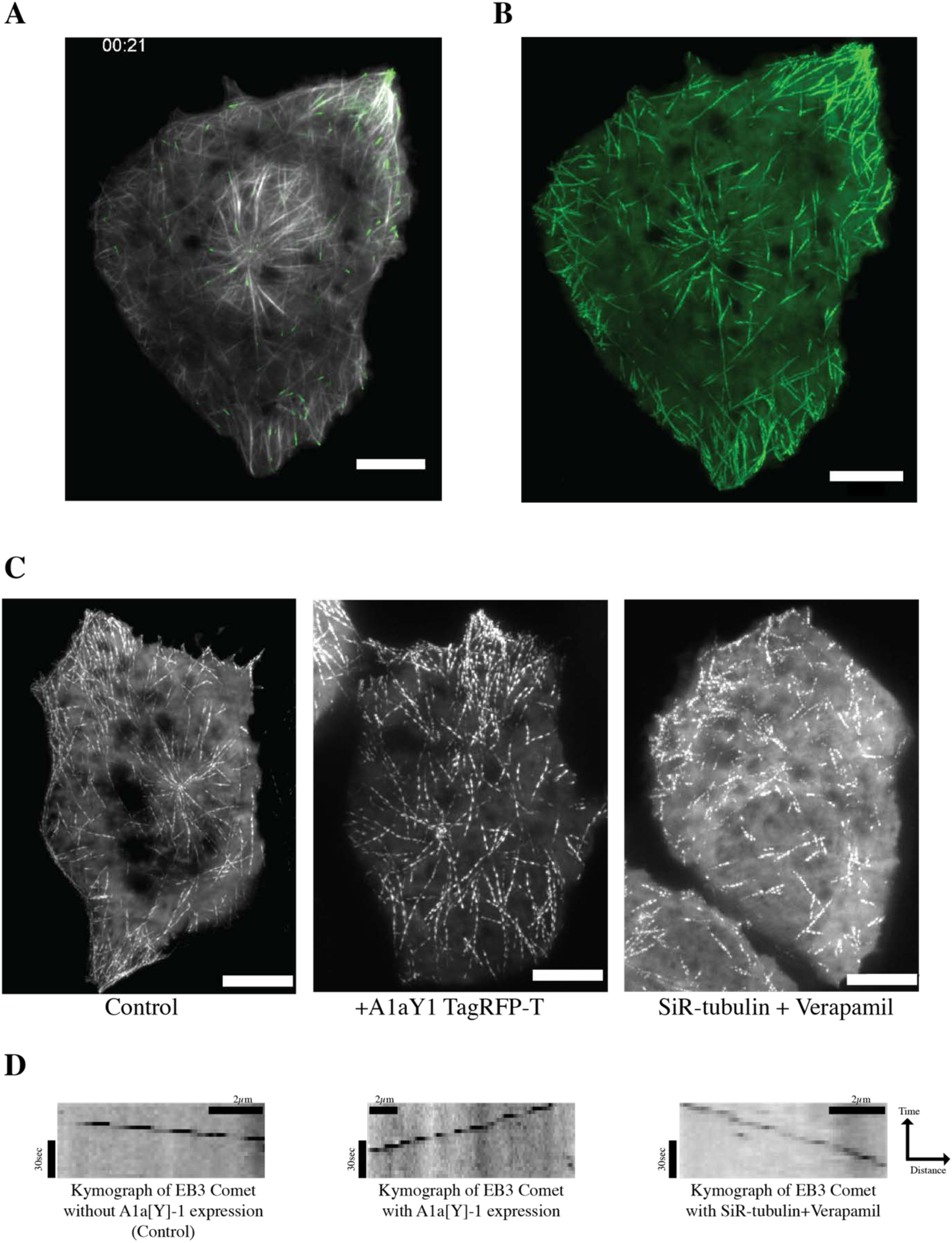
**A.** EB3-GFP (green) marking the growing end of the microtubule decorated with red sensor (grey) stably expressing in U2OS cells. **B.** Z-projection of the first 50 frames of EB3 movement with an interval of 3 sec per frame. **C.** Comparison of Z-projections of EB3 movement for control (without A1aY1 binder), with A1aY1 binder and 0.5μM SiR-tubulin + 10μM Verapamil. Scale bar = 10μm. **D.** Representative kymographs (distance versus time plot) for the growing EB3-GFP comets. Scale bar as indicated.

**Supplementary Figure 8:**
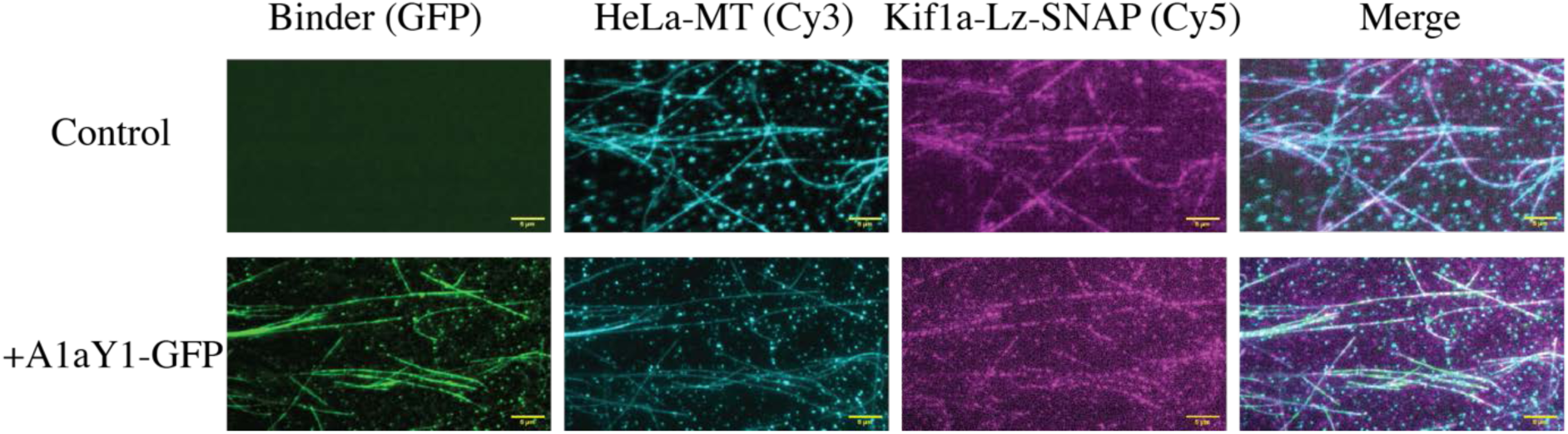
Representative image of single molecule KIF1A motors (shown in magenta) decorated on *in vitro* polymerized HeLa microtubules (shown in cyan) coated with A1aY1-GFP (shown in green). Scale bar = 5μm

## MATERIAL AND METHODS

### Peptides used in screening binders from the library

| Microtubule PTM | Peptide sequence | Residue number on human<br>alpha tubulin | Termed as |
| --- | --- | --- | --- |
| Tyrosination | Biotin-<br>[<sup>440</sup>VEGEGEEEGEEY<sup>451</sup>] | Biotin-TUBA1A 440-451 | *Hs_TUBA1A* |
| Detyrosination | Biotin-<br>[<sup>440</sup>VEGEGEEEGEE<sup>450</sup>] | Biotin-TUBA1A 440-450 | *Hs_TUBA1A-ΔY* |
| Monoglutamylation | Biotin-<br>[<sup>440</sup>VEGEG[E]EEGEEY<sup>451</sup>] | Biotin-TUBA1A 440-451-<br>[E]445 | *Hs_TUBA1A-mG* |

All the above peptide sequences were ordered from Thermo fisher scientific.

### Yeast Surface Display Library Screening

A combinatorial SSO7d yeast display library was obtained as a kind gift from Dr Balaji M. Rao’s lab at Department of Chemical and Biomolecular Engineering, North Carolina State University, Raleigh, NC 27695, USA and the detailed protocol for screening binders was adopted as described in [(Gera et al., 2011)]. To screen a binder for CTT of human alpha tubulin specific for terminal tyrosine residue, SSO7d library was applied for a stringent negative selection with biotinylated detyrosinated (biotin-*Hs*_TUBA1A-ΔY) and mono-glutamylated (biotin-*Hs*_TUBA1A-mG) peptides to reduce the diversity and non-specific binders from the library, followed by a positive selection with tyrosinated peptide (biotin-*Hs*_TUBA1A) yielded a population of binders with varying degree of binding affinities.

The library (diversity around 10^8 cells) was propagated in 10-fold excess of its diversity (10^9 cells) in the fresh glucose-containing SDCAA media [20g/L D-(+)-glucose (Sigma, cat. no. G5767), 6.7g/L yeast nitrogen base without amino acids (BD Difco, cat. no. 291940), 5g/L casamino acids (BD Difco, cat. no. 90001-726), 5.4g/L Na_2_HPO_4_ (Fisher Scientific, cat. no. S374) and 8.6g/L NaH_2_PO_4_.H_2_O (Merck, cat. no. 106346) and 1xPenStrep] thrice at 30°C, 250rpm for 18-24hr before screening. The freshly grown yeast cells were induced (10^9 cells) in galactose containing SGCAA media [20g/L D-(+)-galactose (Sigma, cat. no. G0750), 6.7g/L yeast nitrogen base, 5g/L casamino acids, 5.4g/L Na_2_HPO_4_ and 8.6g/L NaH_2_PO_4_.H_2_O] at 20°C for 20-24hr for the expression of binders on the surface of yeast cells. The freshly induced culture was used for magnetic screening (negative and positive selection with the target peptides) and further sorted using fluorescent activated cell sorter (FACS) to obtain the yeast cells with higher binding affinity for *Hs*_TUBA1A.

#### Magnetic screening

200µl (5*10^6 beads) magnetic beads (Dynabeads Biotin Binder, Invitrogen, Cat. No. 11047) prewashed with PBS-BSA (8g/L NaCl, 0.2g/L KCl, 1.44g/L Na_2_HPO_4_, 0.24g/L KH_2_PO_4_ and 1g/L bovine serum albumin) incubated with 1μM of each biotin tagged peptides (Biotin-TUBA1A 440-451, Biotin-TUBA1A 440-450 and Biotin-TUBA1A 440-451-[E]-445), in separate centrifuge tubes in PBS-BSA at 4°C overnight on a rotatory rod. Around 10^9 (OD600= 1 is around 10^7 cells) freshly induced cells were pelleted and incubated with the 200µl beads coated with biotin-TUBA1A 440-450 and biotin-TUBA1A 440-451-[E]-445 peptide (for negative selection against detyrosinated and monoglutamylated peptides) for 1hr at 4°C on a rotatory rod. Yeast cells bound to the beads were separated using a magnetic rod and discarded. The rest of the unbound cells were incubated for the positive selection with 200µl beads coated with biotin-*Hs*_TUBA1A (tyrosinated peptide biotin-TUBA1A 440-451) at 4°C on a rotatory rod for 1hr. The cells bound to the beads were pulled with the magnetic rod and washed 5 times with PBS-BSA at room temperature for 5-10min. The washed beads were transferred to a fresh 5ml SDCAA media for growth at 30°C, 250rpm for 48hr. Beads were removed from the grown culture after and cells were grown in large culture volume for stocks preparation (10^9 cells) and use for FACS.

#### Fluorescent Activated Cell Sorting (FACS)

The freshly grown cells from positive magnetic screening were induced in galactose containing SGCAA media (10^9 cells) at 20°C for 24hr for surface expression of binder proteins on yeast cells. Around 10^7 cells (OD600= 1) from the culture were pelleted in the microcentrifuge tube. The cells were resuspended in 100µl PBS-BSA and incubated with biotin-TUBA1A 440-451 peptide (100µM) and chicken anti-c-Myc antibody (Invitrogen, Cat. No. A21281) in 1:250 dilution at room temperature for 1hr. Cells were pelleted, washed twice with PBS-BSA and incubated with goat anti-chicken IgG Alexa Fluor 633 (Invitrogen, Cat No. A21052) and Neutravidin fluorescein conjugate (FITC, Invitrogen, Cat. No. A2662, used for 1^st^ FACS) or Streptavidin R phycoerythrin conjugate (Strep-PE, Invitrogen, Cat. No. S866, used for 2^nd^, 3^rd^, 4^th^ FACS) secondary reagent in 1:250 dilutions for 15min on ice. Further cells were washed to remove excess secondary reagents and sorted on BD FACS Aria Fusion for double-positive cells keeping unstained and single stained controls (Central Imaging and Flow Cytometry facility at NCBS). 0.1-1% double-positive cells (for FITC/PE and Alexa fluor-633 fluorophores) were collected to the count of around 4000 to 10,000 cells and grown in 5ml fresh SDCAA media. The freshly grown cells were propagated in larger volumes of SDCAA media (250-500ml) to make stocks and further rounds of sorting experiments. After four rounds of sorting, the cells were plated on an agar plate (20g/L dextrose, 6.7g/L yeast nitrogen base, 5g/L casamino acids, 5.4g/L Na_2_HPO_4_, 8.6g/L NaH_2_PO_4_.H_2_O, 182g/L sorbitol and 15g/L agar) and 10 single yeast colonies were analyzed to identify the most abundant sequence/clone enriched post-screening.

### Protein purification

The gene sequences of A1aY1 and A1aY2 binders were cloned in pET28a(+) vector between NdeI and NotI restriction sites with amino-terminus 6xHis-tag followed by a thrombin site and the gene sequence. The protein expression was achieved in Rosetta (DE3) competent cells by inducing the culture at OD600 of 0.5 with 1mM IPTG (Sigma cat. no. I6758) at 25°C for 8-10hr in terrific broth (Yeast extract 24g/L, Tryptone 20g/L, 17mM KH2PO4, 72mM K2HPO4, 4ml/L Glycerol) and 50µg/ml Kanamycin antibiotic. Only A1aY1 could be purified successfully as A1aY2 protein was toxic to the bacterial expression host (DE3 rosetta) and hence was purified with a carboxy-terminus GFP tag (mentioned below). Overnight-induced culture of A1aY1 was harvested in 50mM Tris-HCl (pH 7.5), 100mM NaCl, 1mM PMSF with 1 tablet of EDTA free protease inhibitor cocktail (Roche, cat. no. 11836170001) for 1L of the culture. The cells were lysed with the Avestin Emulsiflex C3 homogenizer (ATA Scientific instruments) and protein was purified using 5ml Ni-NTA affinity column after 10-column volume (CV) wash with 50mM Tris-HCl (pH 7.5), 500mM NaCl and 25mM Imidazole (pH 7.5) and elution in 5CV of elution buffer (50mM Tris-HCl, pH 7.5, 100mM NaCl and 350mM Imidazole, pH 7.5). The eluted protein was concentrated up to 5ml using 3KDa millipore amicon filter (Merck UFC900324). Further, size exclusion chromatography was carried out in 50mM Tris-HCl (pH 7.5), 100mM NaCl buffer in Superdex-75, 16/600 column (cat. no. 28989333 GE). The resulting fraction of the pure A1aY1 protein was concentrated (mol. wt. 9.3KDa) and frozen in small aliquots for long term storage in −80°C.

For NMR experiments, ^13^C-^15^N isotope-labelled A1aY1 protein was purified from the the DE3 rosetta bacterial cells grown in M9 media (Na2HPO4 6g/L, KH2PO4 3g/L, NaCl 0.5g/L, 15NH4Cl 1g/L, 13C labelled glucose 2g/L, divalent cations, Vit-B12, Thiamine and trace elements and 50µg/ml Kanamycin antibiotic) using Ni-NTA affinity chromatography in Tris buffer as mentioned above. Further purification using size exclusion chromatography was performed in 50mM sodium phosphate buffer (pH 6.5) with 200mM NaCl. Purified protein was set for thrombin cleavage [Thrombin from bovine plasma, Sigma T-4648-10KU (15-20U Thrombin per milligram of the protein)] at room temperature for 4hr. The cleaved protein was purified again with size exclusion chromatography (using S75 16/600 column) in 50mM sodium phosphate buffer (pH 6.5) with 200mM NaCl. The resulting protein corresponds to 7.8KDa which was concentrated and frozen in −80°C in small aliquots for future use in NMR experiments. A GFP tag was attached to the above A1aY1 construct at carboxy-terminus in pET28a(+) vector, such that it contains an amino-terminus 6x-His, a thrombin site, A1aY1 followed by a GFP sequence separated by a glycine-serine linker. Similarly, binder A1aY2 was cloned with a carboxy-terminus GFP tag to obtain its soluble expression. Both the proteins were purified in 50mM potassium phosphate buffer (pH 6.0) with 100mM potassium chloride and 5mM beta-mercaptoethanol (BME) using Ni-NTA chromatography. Further purification of the protein was carried out using S-200, 16/600 column for size exclusion chromatography. For all the *in-vitro* experiments with the polymerized HeLa microtubules, these proteins were diluted in 1xBRB80 (80mM PIPES, 1mM MgCl2, 1mM EGTA, pH 6.8) buffer. For the SPR steady-state binding assay these proteins were diluted in 1xHBS-P+ buffer (10mM HEPES, 150mM NaCl and 0.05% v/v surfactant P20, GE healthcare life science cat. no. BR100671).

#### K560-SNAP and KIF1A-SNAP Purification

Truncated Rat kif1a (1-393 amino acids) followed by a GCN4 leucine zipper was cloned into a pET-17b vector with a Snap-tag followed by a 10X Histidine-tag at the carboxy-terminus. K560-Snap and Kif1a-LZ-Snap were expressed using rosetta (DE3) bacterial expression system. Transformed cells were grown at 37°C to 0.D 0.4-0.6 followed by induction with 0.5mM IPTG (sigma) with overnight shaking at 24°C. Cells were harvested and lysed in buffer A (25mM pipes, pH 6.8, 100mM KCl, 5mM MgCl2, 5mM β-mercaptoethanol, 30mM imidazole). The supernatant was loaded onto a Ni-NTA column, followed by a high salt wash with buffer B (Buffer A with 300mM KCl, 200μM ATP) followed by a high imidazole wash with buffer C (Buffer A with 50mM imidazole) and elution with Buffer E (Buffer A with 350mM imidazole). Pure proteins were obtained by further subjecting the Ni-NTA elute to gel filtration using an S200 16/1600 column (GE) in 1xBRB80. SNAP surface Alexa Fluor 647 (NEB, cat. no. S9136S) labelling of SNAP-tag proteins were performed as per the manufacturer’s protocol on New England BioLabs (NEB) website.

### Cell Culture experiments

Wild type U2OS and HEK293-T cells used in this study were obtained from Prof. Satyajit Mayor’s lab, NCBS Bangalore, India as a gift. U2OS cells were grown in a humidified 37°C incubator with 5% carbon dioxide in McCoy’s 5A (Sigma Aldrich, M4892) media supplemented with 2.2g/l sodium bicarbonate (Sigma, cat. no. S5761), 10% fetal bovine serum (FBS) and 1xPenStrep (cat. no. 15-140-122 Gibco Fisher Scientific). HEK293-T cells were grown in Dulbecco’s modified eagle media (DMEM media from Gibco) supplemented with 1x sodium pyruvate (Gibco), 1xGlutaMAX (Gibco) and 1x PenStrep.

### Mitotic arrest experiment

Stable U2OS cell line expressing A1aY1 with amino-terminus TagRFP-T (red tyrosination sensor) were grown up to 50% confluency in McCoy’s 5A media (Sigma Aldrich, M4892) in 37°C in 5% CO2 incubator. Cells were arrested at S-phase of the cell cycle with 2.5mM thymidine for 16-20hr at 37°C and released for the cell cycle progression for 8-9hr in fresh media. Similarly, a second S-phase arrest in 2.5mM thymidine was performed for 20-24hr and released for 8hr in fresh media, followed by 20ng/ml nocodazole (Sigma Aldrich M1404) treatment for 4hr. The synchronized cells (30-40%) were imaged for mitosis after staining the nucleus of the cells with DAPI (1ug/ml) for 15min or with NucBlue live ready probes reagent (cat. no. R37605 Invitrogen).

### Stable cell line expressing tyrosination sensor

HEK293-T cells were cultured in complete DMEM media with 10% fetal bovine serum, 1xGlutaMAX (Gibco) and 1x PenStrep in humidified 37°C incubator with 5% CO2. Freshly passaged HEK-293T cells were grown up to 70-80% confluency. The gene of interest (A1aY1 cloned with TagRFP-T or TagBFP) was cloned into a lentiviral vector under chicken beta-actin (CAG) promoter flanked with 5’and 3’ LTR sequence. For a 100mm dish transfection, 5µg of lentiviral plasmid cloned with the gene of red or blue tyrosination sensor, 3.75µg of psPAX2 (addgene# 12260) and 1.25µg pmDG2 (addgene# 12259) plasmid were mixed together in 500µl of OptiMEM media with 20µl of P3000 (for Invitrogen Lipofectamine-3000 transfection reagent, Cat. No. L300015) or 10µl of PLUS reagent (for Invitrogen LTX transfection reagent, Cat. No. L15338100). In a separate microcentrifuge-vial took 500µl of OptiMEM and added 30µl of Lipofectamine-3000 or Lipofectamine-LTX reagent. Mixed this Lipofectamine containing solution to the plasmid and incubated at room temperature for 20min. Changed the media of the cells with no PenStrep containing DMEM media and added the transfection mix drop by drop. Cells were incubated for 15-18hr and then changed the media to PenStrep containing media. Following that lentivirus containing supernatant media was collected at 48hr, 72hr and 96hr post-transfection with the replacement of 10ml fresh media every time. The virus supernatant was pooled together and concentrated in 50KDa millipore amicon filter (Merck UFC905024) at 1000*g to around 1-3ml. The concentrated supernatant was supplemented with 1/3 volume of Lenti-X concentrator (Takara Cat. No. 631231) and incubated at 4°C overnight. The viruses were pelleted at 1500*g for 45min at 4°C. The white pellet of lentivirus was resuspended in the 1-2ml of DMEM or McCoy’s complete media (10% FBS) and stored for long term at −80°C in 300-500µl aliquots.

The lentiviral transduction was performed in the 60% confluent culture of wild type U2OS in 5ml complete McCoy’s media (with 10%FBS, 1xPenStrep and 2.2g/l NaHCO_3_) with 1-2μg/ml of polybrene (cat. no. TR-1003-G Merck) and lentiviruses (0.5-1ml thawed at 37°C). After 24hr of transduction, media was changed with the fresh media and cells were propagated as normal cell line stably expressing the red or the blue tyrosination sensor.

### Microtubule tracking experiment (EB3-GFP)

Stable U2OS cell line expressing the red tyrosination sensor (A1aY1 with amino-terminus TagRFP-T) were transiently transfected with 100-500ng of plasmid cloned with end binding protein-3 tagged with GFP (EB3-GFP) using Jet prime transfection reagent (cat. no. 114-15 polypus transfection). EB3-GFP cloned vector was obtained as a gift from Dr Carsten Janke’s lab. Cells were imaged live on Nikon TIRF microscope for the moving EB3-GFP comets with 3sec frame intervals on 37°C stage with 5% carbon dioxide for live-cell imaging. Short movies of 1-5min were acquired and comets were analyzed on FIJI (ImageJ) by generating kymographs (distance vs time) and manually measuring the slope to calculate the velocity of the individual comets. Experiments were performed in triplicates on two different days for control (in wild type U2OS cells without any binder expression), stable U2OS cells (expressing TagRFP-T-A1aY1), and in wild type U2OS cells treated with 0.5µM SiR-tubulin + 10µM Verapamil and imaged post 30min of SiR-tubulin labelling. More than 400 comets were analyzed from 1min movies (3sec frame interval) for all the three sets (450, 468 and 414 comets for control, binder expressing and SiR-tubulin treated cells, respectively) to plot a distribution of the comet velocities from each set. The mean velocities and standard deviations for each set were calculated and distribution of velocities were plotted on Origin lab software.

### Depolymerization assay

Stable U2OS cells expressing red tyrosination sensor (A1aY1 with amino-terminus TagRFP-T) were treated with 10μM nocodazole (Sigma-Aldrich cat. no. M1404) or 0.5mM colchicine (Sigma-Aldrich cat. no. C9754) or 1μM vincristine on a microscope stage set at 37°C with 5% carbon dioxide for live-cell imaging. A time-lapse movie was recorded for microtubule depolymerization event on a Nikon TIRF microscope with 1-3sec frame interval. More than 30min movies were acquired for nocodazole and colchicine treated cells and 5-15min movies were acquired for vincristine treated cells to observe near-complete depolymerisation events of microtubules. Cells were analysed manually for plotting the parameters of polymerisation, depolymerisation, dynamic instability and severing events from the complete frames of the movies. Graphs were plotted on GraphPad Prism6 software.

### Cell viability assay (Trypan blue, propidium iodide (PI) and DAPI staining)

1 million cells of wild type U2OS cell line (as control) and stable U2OS expressing the red or the blue tyrosination sensor were diluted 1000 times in complete McCoy’s media in separate microcentrifuge tubes. 10μl of these 1000 times diluted cultures were mixed with 10μl of 0.4% trypan blue (cat. no. 15250061 Gibco) and the number of dead cells was counted as trypan blue positive cells on an automated cell counter from Invitrogen (supplementary figure6A).

For flow cytometry analysis, PI and DAPI staining were performed on 3-5 million cells of stable U2OS cells expressing the blue tyrosination sensor (A1aY1 with carboxy-terminus TagBFP) and the red tyrosination sensor (A1aY1 with amino-terminus TagRFP-T), respectively (supplementary figure6B). Wild type U2OS cells used as a control in both the staining method. Cells were pelleted at 1000rpm for 5min and resuspended in 2ml 1xPBS (8g/L NaCl, 0.2g/L KCl, 1.44g/L Na_2_HPO_4_, 0.24g/L KH_2_PO_4)_ with 5%FBS. Cells were stained with 1µg/ml concentration of PI and DAPI to mark the dead cells in each culture. For DAPI staining an incubation of 15min was performed on ice. Forward and side scatter was adjusted with blue (488) laser to mark the dense population of the cells for analysis. Percentage viability was calculated by subtracting the number dead cells (DAPI or PI positive cells) from the total number of cells counted in stable U2OS cells with tyrosination sensor and compared it with wild type U2OS control. Cells stained for DAPI and PI were analyzed with violet (405) and green (561) laser, respectively; on BD FACS Aria Fusion cell sorter at CIFF, NCBS Bangalore.

### Transient transfection protocol

All the transient transfection in mammalian cells was carried out at around 60-70% cell confluency in a 35mm Ibidi dishes (Ibidi cat. no. 81156 and 81218 for polymer coverslip and glass-bottom coverslip surface, respectively) using jet prime transfection reagent (cat. no. 114-15 polypus transfection) with 3-5μl of the reagent used per μg of plasmid DNA as per the protocol. For transient transfection of plasmids cloned with the binder (A1aY1) tagged fluorophores (EGFP, mcherry, mCitrine, TagBFP or TagRFP-T) in mammalian cells, 1μg of the respective plasmids were used. Plasmids cloned with detyrosinase VASH2_X1-GFP-2A-SVBP (vasohibin2_X1-GFP separated with 2A sequence followed by small vasohibin binding protein), and polyglutamylases such as eYFP-TTLL5, eYFP-TTLL5 catalytic dead mutant (E366G, ATP deficient mutant), eYFP-TTLL7 were obtained as a kind gift from Dr Carsten Janke’s lab, Institut Curie, France. Around 500ng of these plasmids were used for transfection in stable U2OS cells expressing the red tyrosination sensor (A1aY1 with amino-terminus TagRFP-T). All the transfections were performed in 10% serum-containing media and cells were transfected for 4-8hr, followed by which the media was changed with fresh complete media. Cells were stained with SiR-tubulin (0.5-1μM, cat. no. CY-SC002 Spirochrome kit, Cytoskeleton, Inc) for 1hr before imaging. Cells were imaged on FV3000 Olympus confocal microscope post 24hr of transfection.

### SPR (Surface Plasmon Resonance) steady-state binding assays

All the binding assays were performed on Biacore-T200 instrument from GE Healthcare Life Sciences. GE SA sensor chip (cat. no. BR100531) was immobilized with 1μg/ml concentrations of the following biotin peptides; biotin-TUBA1A 440-451 (biotin-*Hs*_TUBA1A), biotin-TUBA1A 440-450 (biotin-*Hs*_TUBA1A-ΔY), biotin-TUBA1A 440-451-[E]-445 (biotin-*Hs*_TUBA1A-mG), biotin-Tub84B 438-450 (biotin-*Dm*_Tub84B), biotin-TBA1 437-349 (biotin-*Ce*_TBA1). A single SA chip can be immobilized with three different peptides at flow channel (FC) 2, 3 and 4 keeping the FC1 as blank for buffer. Two SA sensor chips were used to perform assays with the five peptides mentioned above keeping biotin-*Hs*_TUBA1A as common. The first chip was immobilized with biotin-*Hs*_TUBA1A, biotin-*Hs*_TUBA1A-ΔY, and biotin- *Hs*_TUBA1A-mG peptides; while the second chip was immobilized with biotin- *Hs*_TUBA1A, biotin-*Dm*_Tub84B, and biotin-*Ce*_TBA1 peptides. The surface of the SA chip was pre-activated with NaCl and NaOH solutions (as per the manufacturer’s protocol in GE manual for SA surface immobilization) before peptide immobilization. All the peptides were immobilized in the range of 100-300 response unit (RU). All the assays were performed at 30μl/min flowrate in the 1xHBS-P+ buffer (10mM HEPES, 150mM NaCl and 0.05% v/v surfactant P20, GE healthcare life science cat. no. BR100671) in triplicates with two different batches of the protein. The GFP tagged binders (A1aY1 and A1aY2) were tested for their binding parameters (association and dissociation) by titrating 8μM of both the proteins (A1aY1-GFP and A1aY2-GFP) on the chip immobilized with biotin-*Hs*_TUBA1A peptide (Biotin-TUBA1A 440-451) and plotted the relative response (RU) for 180sec contact time, 300sec dissociation time, 2 regeneration steps with 10mM Glycine-HCl (each time with pH2.5) for 30sec. With the sensogram obtained in supplementary figure-1C showed positive binding with A1aY1-GFP while no apparent binding with A1aY2-GFP.

To determine the binding interaction (k_d_) of A1aY1, a steady-state binding assay (considering 1:1 binding) with a range of different concentrations (0.031μM, 0.062μM, 0.125μM, 0.25μM, 0.5μM, 1μM, 2μM, 4μM, 8μM, 15μM, 30μM, 50μM, 100μM and 200μM) were titrated to all the immobilized peptides on FC2, FC3 and FC4 of SA chip with 120sec contact time, 180sec dissociation, 2 regeneration step of 30sec each with 10mM Glycine-HCl pH 2.0 and 2.5 respectively followed by 60sec stabilization period between two titrations. The relative responses of binding (RUmax) with each peptide were determined and normalised (maximum response as 100) by subtracting the blank flow channel 1 (as FC2-1, FC3-1 and FC4-1), and was plotted with the corresponding concentrations to fit a curve (one site total fitting on GraphPad Prism6) and determine the value of Kd.

### Alpha tubulin C-terminal tail (CTT) peptides

*Hs*_TUBA1A, *Hs*_TUBA1A-ΔY and *Hs*_TUBA1A-mG peptides were synthesized from ThermoFisher and were used in screening (as mentioned above) and SPR SA chip immobilization. *Drosophila melanogaster* and *C. elegans* alpha-tubulin CTT peptides; biotin-Tub84B 438-450 (biotin-*Dm*_Tub84B; biotin-^438^SGDGEGEGAEEY^450^) and biotin-Tba-1 437-449 (biotin-*Ce*_TBA1; biotin-^437^SNEGGNEEEGEEY^449^) were synthesized from Lifetein and were used in SPR SA chip immobilization.

### NMR Spectroscopy

Protein samples were prepared as described above. All NMR spectra were acquired at 25°C on an 800MHz/600MHz Bruker Avance III spectrometers equipped with a 5mm TCI CryoProbe. The sample was loaded in 5mm Shigemi tube. ^1^H–^15^N heteronuclear single quantum coherence (HSQC) experiments were performed with 2048 × 256 complex data points. All NMR spectra were processed by using NMRPipe***^[1]^*** and analyzed by using Sparky***^[2]^***.

Assignment of the backbone resonances (^1^H, ^13^C, and ^15^N) of the proteins were carried out by using Bruker’s BEST pulse program (b_HNCO, b_HNCACO, b_HNCACB, and b_CBCACONH) experiment. ^1^H and ^13^C resonance assignments of side-chain atoms in A1aY1 were obtained by collecting H(CC)CONH (H)CC(CO)NH and HCCH-TOCSY respectively. The resonance list for the above experiments generated by using sparky and submitted to I-PINENMR***^[3]^*** server for automatic assignment. The I-PINE results are manually cross-checked and corrected. The model structure of A1aY1 was calculated by using CS-ROSETTA***^[4]^***. The interface between A1aY1 and nonbiotinylated *Hs*_TUBA1A peptide was identified using ^13^C filtered NOESY experiments on a 1:1 sample of ^13^C, ^15^N enriched protein, and unlabeled (natural abundance) Peptide.

NMR titration was performed by titrating nonbiotinylated *Hs*_TUBA1A and *Hs*_TUBA1A-ΔY peptides (ligand) to the 300μM sample of ^13^Cand ^15^N-labeled A1aY1 (protein) on Bruker Ascend 600 MHz spectrometers equipped with triple resonance cryoprobes and pulsed-field gradients. ^1^H-^15^N HSQC spectra of each residue were taken for all the titrations. For each titration point (typically 0.5, 1, 2, 3, 4, 5, 6, 7 equivalents of ligand) a two-dimensional water-flip-back ^15^N-edited HSQC spectrum was acquired with 2048 (256) complex points, 100ms (60ms) acquisition times, apodized by 60 shifted squared (sine) window functions and zero-filled to 1024 (512) points for ^1^H and ^15^N, respectively. Chemical shift perturbations (CSP) were calculated for individual amino acids in the ^13^C and ^15^N-labeled A1aY1 protein from the saturating titration (1:7 molar excess of the protein:peptide) with the tyrosinated *Hs*_TUBA1A peptide (supplementary figure2D). For each residue, the weighted average of the ^1^H and ^15^N CSP was calculated as CSP = [(δHN_b_^-^δHN_f_)^2^ + (δN_b_^-^δN_f_)^2^/25)]^1/2^, where δHN_b_and δHN_f_ are the ^1^HN amide chemical shifts of bound and free form, respectively. Similarly, the δN_b_ and δN_f_ are the ^15^N amide chemical shifts of bound and free form, respectively.

### In-vitro motor gliding assay on HeLa microtubules

HeLa tubulins were obtained from Dr Carsten Janke’s lab, Institut Curie, France. 10μM tubulin was set for polymerization in 1xBRB80 buffer (80mM PIPES, 1mM MgCl_2_, 1mM EGTA, pH 6.8 with KOH) in presence of biotin-labeled tubulin (1/10 molar ratio), Alexa-fluor-561 labeled tubulin (1/10 molar ratio), 2mMGTP (Sigma) and 20μM taxol (Sigma) at 37°C. A glass chamber made up with coverslip glass (0.17mm thick) was passivated with BSA-biotin (Thermo Scientific, cat. no. 29130, 1mg/ml, 5min) followed by a 1xBRB80 wash and 0.5mg/ml streptavidin (Thermo Scientific, cat. no. 43-4302) coating for 5min. The surface was blocked with 5% Pluronic-F127 (Sigma, cat. no. P2443) and 1.25mg/ml beta-casein (Sigma, cat. no. C6905) in 1xBRB80 buffer. 10-20 fold diluted polymerized HeLa microtubules were flowed in the flow chamber and then washed with 1xBRB80 containing 0.05% Pluronic-F125, 1.25mg/ml beta-casein and 20μM Taxol. The binder (50μg/ml A1aY1-GFP) and single-molecule (nanomolar) concentration of the motors (K560-SNAP and KIF1A-LZ-SNAP labelled with 640 SNAP-tag) had flowed in the flow chamber and imaged the flow chamber with blue (488) laser, green laser (561) and red laser (640) for binder, HeLa microtubule and single molecule motor movement, respectively on Nikon H-TIRF microscope.

### 3D-SIM (Structural Illumination Microscopy)

#### 3D-SIM acquisition

U2OS cells stably expressing the red tyrosination sensor (TagRFP-T-A1aY1) were plated on glass-bottom (#1.5H) ibidi dishes and examined using Nikon N-SIM fitted with 100x/1.49 SR Plan Apo TIRF oil immersion objective. Image stacks (z-steps of 0.2μm) were acquired with Andor iXon3 (DU-897) EM-CCD camera. Exposure conditions were adjusted to get a typical yield of about 3000 max counts while keeping the bleaching minimal. Image acquisition, SIM image reconstruction and data alignment were performed using NIS-Elements 4.2 software (Nikon). Image reconstruction was done by varying image modulation contrast @ 0.5-1.0, High-resolution noise suppression @0.50-1.0, Out of focus blur suppression @0.1-0.2.

#### Full-Width Half Maximum (FWHM) estimation

Line profiles of several microtubules from 10 labelled cells were obtained and fitted with a Gaussian distribution using Fiji image processing software. For the profile, a line width of 165 nm was chosen to cover a minimum width of four pixels in SIM images. Microtubule widths in SIM stacks were measured in the z-section where the microtubule fluoresced most strongly. The full width at half maximum (FWHM) was calculated with the following formula where σ is the Gaussian width parameter. FWHM = 2.355σ

### Cell fixation, immunofluorescence and imaging

HeLa cells plated on glass coverslips were transduced with lentiviruses encoding different constructs of mVash-mSVBP for 24 hours and were fixed according to previously described protocols**^5^**. Briefly, the cellular proteins were cross-linked using the homo-bifunctional cross-linker, Dithio-bis(succinimidyl propionate; DSP; #22585 Thermo Fisher Scientific) diluted in microtubule-stabilizing buffer (MTSB), followed by fixation with 4% paraformaldehyde for 15 mins. The cells were then permeabilized with 0.5% Triton X-100 in MTSB for 5 mins and were blocked in 5% bovine serum albumin (BSA) prepared in PBS containing 0.1% Triton X-100 (PBST).

Cells were incubated with primary antibodies (Anti-tyrosinated tubulin antibody YL1/2, 1/5,000; Abcam #ab6160, Anti-detyrosinated tubulin antibody, 1/1,000; Merck #AB3201, Anti-α-tubulin antibody 12G10, 1/1,000, developed by J. Frankel and M. Nelson, obtained from the Developmental Studies Hybridoma Bank, developed under the auspices of the NICHD, and maintained by the University of Iowa)) diluted in PBST containing 5% BSA (blocking solution) for 2 h at room temperature. Cells were then incubated with secondary antibodies (goat anti-rabbit Alexa Fluor 568, 1:1,000; Thermo Fisher Scientific #A11036, goat anti-rat Alexa Fluor 594, 1:1,000; Thermo Fisher Scientific #A-11007, goat anti-mouse Alexa Fluor 568, 1:5,000; Thermo Fisher Scientific #A11019) prepared in blocking solution and incubated for 1 h at room temperature. Coverslips were mounted using ProLong Gold anti-fade medium (#P36930, Thermo Fisher Scientific).

Images were acquired using Optigrid (Leica systems) with a 63x (numerical aperture 1.40) oil immersion objective and the ORCA-Flash4.0 camera (Hamamatsu), operated through Leica MM AF imaging software. Images were processed using ImageJ v1.51a (National Institute of Health) and final figures were prepared using adobe photoshop and illustrator.

### Sample preparation and Immunoblotting

HeLa cells transduced with lentiviruses encoding for different constructs of mVash-mSVBP for 48 hours were directly collected in 2x Laemmli buffer (180 mM DTT (Sigma #D9779), 4% SDS (VWR #442444H), 160 mM Tris-HCl pH 6.8, 20% glycerol (VWR #24388.295), bromophenol blue). The samples were then boiled at 95°C for 5 min, spun down at 20,000×g for 5 min using a table top centrifuge, and stored at −20°C. Immunoblotting was performed according to previously described protocols***^[5]^***. Briefly, SDS-PAGE gels were prepared at 375 mM Tris-HCl, pH 9.0, 0.1% SDS (Sigma Aldrich #L5750) and 10% acrylamide (40% acrylamide solution (Bio-Rad #161-0140) supplemented with 0.54% bis-acrylamide (w/v) powder (Bio-Rad #161-0210)). Samples were loaded on SDS-PAGE gels, separated and transferred onto a nitrocellulose membrane (Bio-Rad #1704159) using Biorad Trans-Blot^®^ Turbo system, according to manufacturer’s instructions. Membranes were blocked for 1 h in 5% non-fat milk prepared in phosphate-buffered saline (PBS) containing 0.1% Tween-20 (PBS-T). After blocking, membranes were incubated with primary antibodies (Anti-tyrosinated tubulin antibody YL1/2, 1/5,000; Abcam #ab6160, Anti-detyrosinated tubulin antibody, 1/5,000; Merck #AB3201, polyclonal rabbit anti-GFP antibody (1:5,000; Torrey Pines Biolabs TP401, Anti-α-tubulin antibody 12G10, 1/1,000, developed by J. Frankel and M. Nelson, obtained from the Developmental Studies Hybridoma Bank, developed under the auspices of the NICHD, and maintained by the University of Iowa) for prepared in PBS-T containing 2.5% non-fat milk for 2 h. Membranes were washed 4x with PBS-T, and then incubated with HRP-conjugated secondary antibodies (goat anti-rabbit, 1:10,000; Bethyl #A120-201P, goat anti-mouse, 1:10,000; Bethyl #A90-516P, goat anti-rat, 1:10,000; Bethyl #A110-236P) for 1 h. Membranes were then washed 4x in PBS-T and Chemiluminescence signal on the membrane was revealed using Clarity™ Western ECL substrate (Biorad #1705060) solution.

### DNA constructs and lentivirus production for Vashohibin and SVBP

mVash and mSVBP were cloned in to pTRIP lentiviral vectors using sequence- and ligation-independent cloning***^[6]^***, which was described in detail previously***^[7]^***. Briefly, mSVBP was amplified from cDNA prepared from cultured hippocampal neurons using primers with at least 15 bp of homology sequence to the ends of the pTRIP vector with CMV-enhanced chicken beta-actin (CAG) promoter at the XhoI site. mSVBP was cloned after a self-cleavable 2A peptide sequence***^[8]^*** which was in frame with the GFP. mVash2_X3 was amplified from brain cDNA and cloned into pTRIP-mSVBP vector at the NheI site to generate mVash2_X3-2A-mSVBP. mVash1 and mVash2_X1 were amplified from testis cDNA and cloned into pTRIP-mSVBP vector at the NheI site to generate mVash1-2A-mSVBP and mVash2_X1-2A-mSVBP, respectively. Primers used for amplification are given below: mSvbp-FS-2A: 5’-tccactagtgtcgac ATGACTACTGTCCCATTGTGCAGG-3’, mSvbp-RS-2A: 5’-ttttctaggtctcgag TtACTCCCCAGGCGGCTGCATCTG-3’, mVash2-FS1: 5’-agaattattccgctagc ATGACCGGCTCTGCCGCCGACAC-3’, mVash2-RS1: 5’-accatGGTGGCgctagcGATCCGGATCTGATAGCCCACTTCG-3’, mVash1-FS1: 5’-agaattattccgctagc ATGCCAGGGGGAAAGAAGGTGGTC-3’ and mVash1-RS1: 5’-accatGGTGGCgctagc CACCCGGATCTGGTACCCACTGAG-3’. Packaging plasmids psPAX2 and pCMV-VSVG are gifts from D. Trono (Addgene plasmid #12260) and B. Weinberg (Addgene plasmid #8454), respectively.

Lentivirus particles were produced using protocols described earlier(Bodakuntla et al., 2019). Briefly, X-Lenti 293T cells (#632180; Takara) were co-transfected with 1.6µg plasmid of interest and packaging plasmids (0.4µg of pCMV-VSVG and 1.6µg of psPAX2) using 8µl of TransIT-293 (#MIR 2705, Mirus Bio) transfection reagent per well of a 6-well plate. Next day, the culture medium of the X-lenti cells was changed to Neurobasal medium (Thermo Fisher #21103049) containing 1× Penicillin-Streptomycin (Life Technologies #15140130). After 24 – 30hr of media change, the virus-containing supernatants were passed through a 0.45µm filter and either used fresh or aliquoted and stored at −80°C. To determine the optimal amount of virus for transduction, lentivirus aliquots were thawed, and different volumes were added to HeLa cells. Based on the expression levels of the GFP protein, the desired amount of virus to achieve maximum transduction efficiency was determined.

### Image acquisition and analysis

All the confocal imaging was performed on Olympus FV3000 inverted microscope equipped with 60x oil objective (1.42 NA) and six solid-state laser lines (405, 445, 488, 514, 561, and 640). All the images were acquired in 2048*2048 frame (at 60x objective) marking the region of interest for acquisition. Optical sections of 0.5-1μm (step size) are z-projected in all image panels shown in figure2, figure3, supplementary figure3, 4 and 5. All the images acquired were analyzed on FIJI software (ImageJ) using appropriate plugins.

Pearson’s correlation coefficient (R-value) calculation in figure-3B was carried on z-projections of the images acquired on FV3000 confocal microscope for each enzyme overexpression. For each enzyme experiment, the confocal stacks were z-projected for TagRFP-T (561 laser channel) and SiR-tubulin channel (640 laser channel). A 50-pixel background was subtracted from both the channel. The colocalisation was calculated using ImageJ coloc-2 plugin. Pearson’s R-value (no threshold) was calculated for each image, which represents the percentage overlap of the fluorescent intensity TagRFP-T over SiR-tubulin.

A single plane imaging for microtubule growth rate (EB3 experiments), motor movement and microtubule drug-based depolymerization were performed on Nikon Ti2 H-TIRF microscope with 100x oil objective (1.49 NA) having four solid-state lasers (405, 488, 561 and 640) under total internal reflection mode. The images were acquired using an appropriate filter set using the s-CMOS camera (Hammamatsu Orca Flash 4.0) controlled by Nikon NIS-elements software. All the images were analyzed on FIJI (ImageJ) software to construct movies and kymographs to measure the velocities of motor and EB3 comets.

### Graphs and statistical analysis

All the graphs and statistical analysis were carried out on Origin lab (student version) and commercially available GraphPad Prism6 software. The SPR steady-state binding graph in figure-1B, line scan intensity profile in figure-3A, Pearson’s coefficient graph in figure-3B, sensogram of supplementary figure-1C, CSP plot in supplementary figure-2D and graphs in figure-5 were plotted on GraphPad Prism6 software. Graphs in figure-4 for EB3-GFP comet and motor velocities were plotted for histograms and Gaussian fit on Origin lab software.

